# Structural interpretation of the effects of threo-nucleotides on nonenzymatic template-directed polymerization

**DOI:** 10.1101/2020.11.17.387142

**Authors:** Wen Zhang, Seohyun Chris Kim, Chun Pong Tam, Victor S. Lelyveld, Saikat Bala, John C. Chaput, Jack W. Szostak

**Author notes:** Beam Therapeutics, 26 Landsdowne St., Cambridge, MA 02139, USA. Polypeptide Laboratories Inc., Torrance, CA 90503, USA.

## Abstract

The prebiotic synthesis of ribonucleotides is likely to have been accompanied by the synthesis of noncanonical nucleotides including the threo-nucleotide building blocks of TNA. Here we examine the ability of activated threo-nucleotides to participate in nonenzymatic template-directed polymerization. We find that primer extension by multiple sequential threo-nucleotide monomers is strongly disfavored relative to ribo-nucleotides. Kinetic, NMR and crystallographic studies suggest that this is due in part to the slow formation of the imidazolium-bridged TNA dinucleotide intermediate in primer extension, and in part because of the greater distance between the attacking RNA primer 3’-hydroxyl and the phosphate of the incoming threo-nucleotide intermediate. Even a single activated threo-nucleotide in the presence of an activated downstream RNA oligonucleotide is added to the primer ten-fold more slowly than an activated ribonucleotide. In contrast, a single activated threo-nucleotide at the end of an RNA primer or in an RNA template results in only a modest decrease in the rate of primer extension, consistent with the minor and local structural distortions revealed by crystal structures. Our results are consistent with a model in which heterogeneous primordial oligonucleotides would, through cycles of replication, have given rise to increasingly homogeneous RNA strands.

## INTRODUCTION

Studies of the path from the prebiotic world to the RNA World have long contended with the chemical complexity of RNA. Eschenmoser proposed a comprehensive synthetic approach, suggesting that the origin of RNA might be elucidated by a systematic exploration of the properties of its chemical analogues and structural relatives(1,2). Such a constructive approach addresses several seemingly intractable questions in abiogenesis. In particular, are ribonucleic acids chemically unique in their informational and functional properties? Is it possible that a chemically simpler genetic polymer might have arisen prior to RNA? Or, could RNA have been the winner in a competition with alternative polymers? It was quickly recognized that, in order to have been a true ancestor of RNA, any such a polymer would require the ability to transmit its information to RNA(3). This broad pursuit might be further constrained by recognizing that, to be a viable genetic polymer preceding or competing with RNA, such a species must also be capable of being replicated in a template-directed manner by a nonenzymatic process.

Eschenmoser and colleagues proposed and synthesized a promising candidate in this search: threose nucleic acid (TNA). The TNA backbone is composed of phosphodiester-linked 3’➔2’ α-L-threofuranose sugars and, therefore, has one fewer atom in its repeating unit than RNA(4). The possible prebiotic relevance of TNA has been bolstered by the observation that threose is one of the higher yield products of Kiliani-Fischer sugar synthesis(5). Threose is chemically simpler than ribose not only because of its shorter carbon chain length but also its reduced stereochemical complexity (Figure 1A,B). Prebiotically plausible pathways for producing threo-nucleotides have recently been proposed(6). TNA synthesis is simpler than RNA synthesis, because the threose sugar component of TNA can be generated from two molecules of glycolaldehyde, whereas the ribose component of RNA requires the sequential reaction of glycolaldehyde followed by glyceraldehyde. Critically, TNA can form stable Watson-Crick duplexes with itself, DNA, and RNA, such that the exchange of genetic information between these polymers appears feasible (Figure 1C)(4). Structural studies demonstrate that TNA binds RNA to form right-handed double-helical duplexes(7–9) despite the fact that the contracted chemical structure of the TNA backbone decreases the distance between sequential phosphorus atoms.

**Figure 1.**
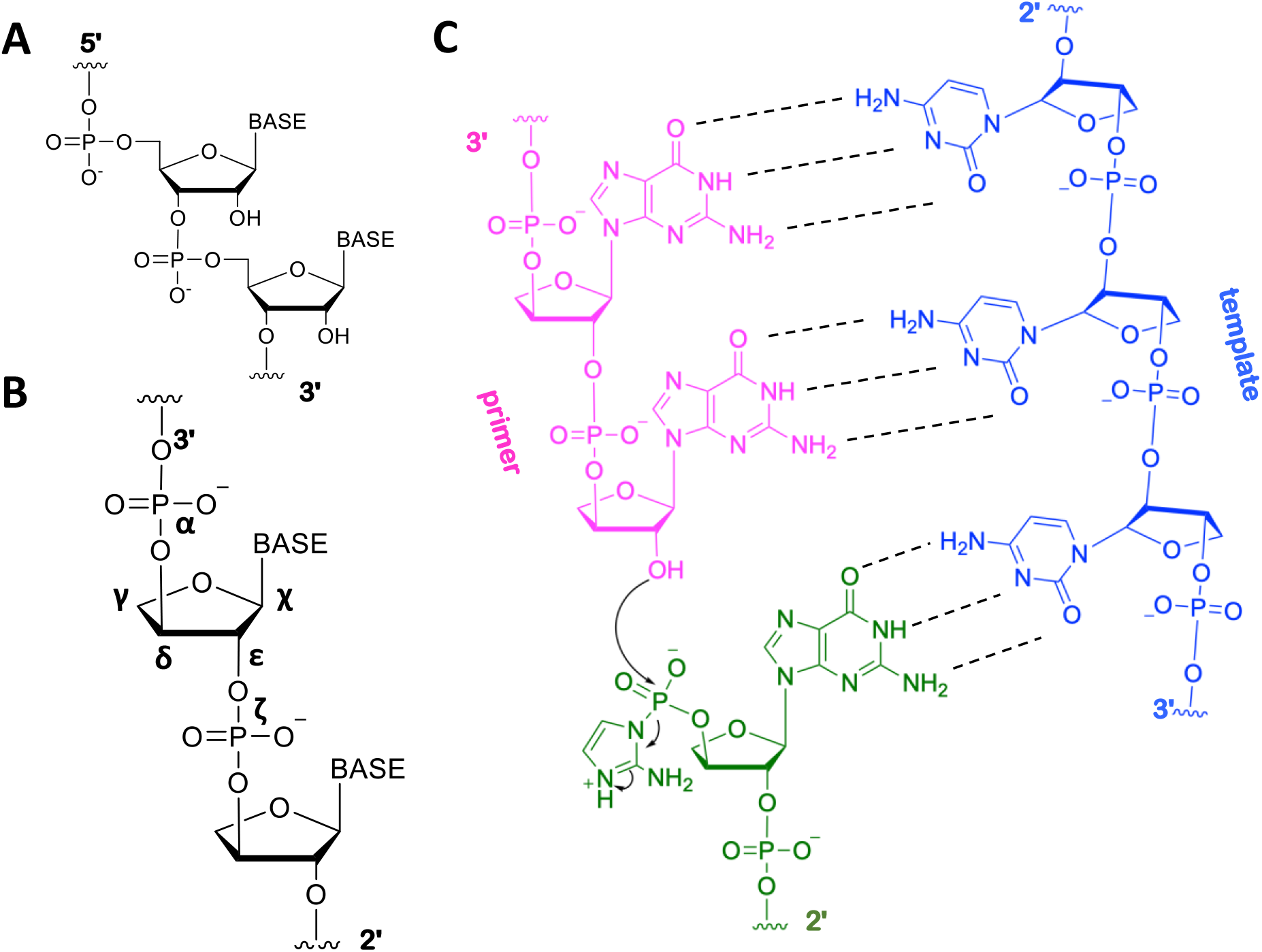
Comparison of RNA and threose nucleic acid (TNA) structures. (A) Native RNA chemical structure. (B) TNA chemical structure. Backbone and glycosidic torsion angles are labelled. (C) Chemical representation of hypothetical TNA template (blue) directed ligation of a TNA primer (pink) and a TNA ligator (green), of which the 3’-phosphate is activated as a 2-aminoimidazolide.

To address the question of the replicative capacity of TNA, we have previously reported the enzymatic copying of DNA to TNA and vice versa, using both wild type and engineered polymerases(10–12). However, it remains unclear whether TNA can support efficient template-dependent copying by nonenzymatic chemistry. Heuberger and Switzer have demonstrated nonenzymatic synthesis of guanosine oligo-ribonucleotides on a TNA template(13). We subsequently reported nonenzymatic template-directed primer extension using 2-methylimidazole-activated 2’-amino-threose nucleotides as substrates, using DNA, RNA and TNA templates(14). In both cases, however, the TNA-templated nonenzymatic copying reactions were much slower than the corresponding RNA-templated reactions, irrespective of the precise identity of the activated mononucleotides used in primer extension reactions. More recently the Krishnamurthy group showed that oligomers of amino-terminated TNA or RNA oligonucleotides could be ligated together with similar efficiency on all-RNA templates, but that ligation on chimeric TNA-RNA templates was much less efficient(15). The consistent observation of slow kinetics for TNA-directed nonenzymatic copying provides a biophysical argument for the emergence of RNA as opposed to TNA based life.

Here we examine the behaviour of TNA in nonenzymatic template-directed primer extension reactions from both a kinetic and a structural perspective. We find that activated TNA monomers alone are not effective substrates for multi-step nonenzymatic primer extension reactions, and even single activated threo-nucleotides in an otherwise all RNA context are incorporated with poor efficiency. Through a series of kinetic and co-crystallization studies, we analyse the origins of these experimental observations. Our results suggest that the reactive intermediate in primer extension, an imidazolium-bridged dinucleotide composed of 3’-3’ linked threo-nucleotides, is both generated very slowly and once formed, binds to the template with suboptimal geometry for primer 3’-OH attack. In contrast we find that a single TNA nucleotide at the primer terminus or at the +1 template site only modestly slows the rate of primer extension, consistent with crystallographic observations that such substitutions have relatively minor structural consequences. Taken together, we propose that structural constraints explain the diminished ability of TNA to engage in nonenzymatic template-directed primer extension, and we discuss the consequences of these observations for the plausibility of the involvement of TNA in the origin of primitive cells.

## MATERIALS AND METHODS

### RNA and TNA oligonucleotides preparation

Native, LNA-containing, and TNA-containing RNA oligonucleotides used for primer extension and crystallographic studies were custom-synthesized by Exiqon Inc. (Woburn, MA) or IDT Inc. (San Jose, CA), or were synthesized in-house on an Expedite 8900 DNA/RNA synthesizer. TNA phosphoramidites were synthesized as previously reported(16,17). Oligonucleotides synthesized in-house were deprotected using AMA (1:1 v/v aqueous mixture of 30% w/v ammonium hydroxide and 40% w/v methylamine) for 20 min at 65 °C, followed by desilylation with Et_3_N·3HF. Oligonucleotides were HPLC purified on an Agilent ZORBAX Eclipse-XDB C18 column using 25 mM triethylammonium bicarbonate in H_2_O (pH 7.5) with gradient elution from 0 % to 30 % acetonitrile over 40 mins. The oligonucleotides were collected, lyophilized, desalted, and concentrated as appropriate for primer extension and crystallization experiments. Oligonucleotides were characterized by LC-MS (see SI for details).

### Synthesis of α-L-threofuranosyl-3’-phosphoro-(2-aminoimidazolide) (2-AIptG and 2AIptC)

The activated monomers 2-AIptG and 2AIptC were synthesized essentially as previously reported for the corresponding activated ribonucleotides(18). Briefly, to a solution of 30 mM α-L-threofuranosyl-3’-monophosphate (tGMP and tCMP)(17,19) and 2-aminoimidazolium chloride (5 eq.) in DMSO was added triphenylphosphine (PPh_3_, 10 eq.), triethylamine (TEA, 5 eq.) and 2,2’-dipyridyldisulfide (DPDS, 9 eq.) under stirring at room temperature. The reaction was allowed to proceed for 4 h, followed by precipitation of the crude product, supernatant removal and product purification by reverse phase flash chromatography. For detailed experimental procedures, see Supporting Information 1.4.

### Primer extension assays

Primer (1 μM) and template (1.5 μM) were pre-mixed in 15 μL of a solution containing 200 mM HEPES pH 8.0 and 200 mM MgCl_2_. The primer extension reaction was initiated by adding the indicated activated monomers (2-AIptG, 2-AIptC, 2-AIpC, or 2-AIpG) from 200 mM stock solutions to a final concentration of 40 mM in the reaction. At the indicated time points, 1 μL aliquots were removed and quenched with 29 μL of 90 % (v/v) aqueous formamide containing 15 mM EDTA. Samples were heated at 95 °C for 2 min, cooled to room temperature, and analysed on 18 % (19:1) denaturing PAGE gels containing 7 M urea. Gels were imaged using a Typhoon 9410 scanner. The resulting gel images were analysed using ImageJ software.

### Crystallization

RNA samples (0.5 mM duplex or hairpin in water) were heated to 80 °C for 5 min, slowly cooled to room temperature (20 °C), and incubated at 4 °C overnight prior to crystallization. Nucleic Acid Mini Screen Kits, Natrix (Hampton Research) and Nuc-Pro-HTS (Jena Bioscience) were used to screen crystallization conditions at 18 °C using the sitting-drop method. An NT8 robotic system and Rock Imager (Formulatrix, Waltham, MA) were used for crystallization screening and monitoring the crystallization process. Optimal crystallization conditions are listed in the Supporting Information.

### Data collection, structure determination and refinement

Crystals were mounted and soaked for 2 min in a drop of mother liquor solution which contained 30% glycerol as a cryo-protectant, then transferred to liquid nitrogen. Diffraction data were collected at wavelength of 1 Å under a liquid nitrogen stream at −174 °C on Beam line 821 or 822 at the Advanced Light Source in the Lawrence Berkeley National Laboratory (USA). The crystals were exposed for 1 s per image with a 1 ° oscillation angle. The distances between detector and the crystal were set to 200 - 300 mm. Raw diffraction data was processed using HKL2000(20). Structures were solved by molecular replacement(21) using our previously determined structures (PDB codes 5VCI, 5HBX or 5HBW) as search models. All structures were refined using Refmac(22) in the CCP4 suite. The refinement protocol included simulated annealing, positional refinement, restrained B-factor refinement, and bulk solvent correction. The topologies and parameters for locked nucleic acids, threose mononucleotide (tGMP), threose nucleotide residues (tA and tG) and 3’-3’-imidazolium-bridged di-threose guanosine intermediate (tGp-AI-ptG) were created using ProDrg in the CCP4 suite(23) and applied during refinement. After several cycles of refinement, a number of ordered waters were added. Data collection, phasing, and refinement statistics of the determined structures are listed in the Supporting Information.

## RESULTS

### Activated threo-nucleotides are poor substrates for nonenzymatic primer extension

#### Primer extension using 2-AIptG as substrate

We first sought to determine whether an RNA primer could be extended with multiple activated threo-nucleotide monomers. To generate the monomer, we activated α-L-threofuranosyl guanosine 3’-monophosphate (tGMP) to its corresponding 3’-phosphoro-2-aminoimidazolide (2AIptG) following the standard procedure for the preparation of activated ribo-nucleotides(24) (details in Supporting Information). For our initial test of primer extension using activated tG, we used an all-RNA primer and an RNA template oligonucleotide with the templating sequence 3’-CCCCAA-5’ (Figure 2)(16). Although primer extension was observable in the presence of 40 mM activated threo-nucleotide monomer 2AlptG, it was extremely slow (est. less than 0.1 h^-1^). In contrast, primer extension reactions containing the activated ribo-nucleotide monomer 2-AIpG yielded >95% extended product within one hour, and the observed rate constant for disappearance of unreacted primer was 12 h^-1^ (Figure 2B). Thus primer extension with 2AIptG alone is at least 100-fold slower than with 2AIpG, in the context of an RNA primer and an RNA template. We considered the possibility that the primer template context might affect primer extension with threo-nucleotides, e.g. geometrical constraints might lead to a requirement for a TNA primer and/or a TNA template. However, all tested combinations including a primer ending in a TNA residue and a TNA template led to even slower primer extension with 2AIptG as the monomer (Figure S-1).

**Figure 2.**
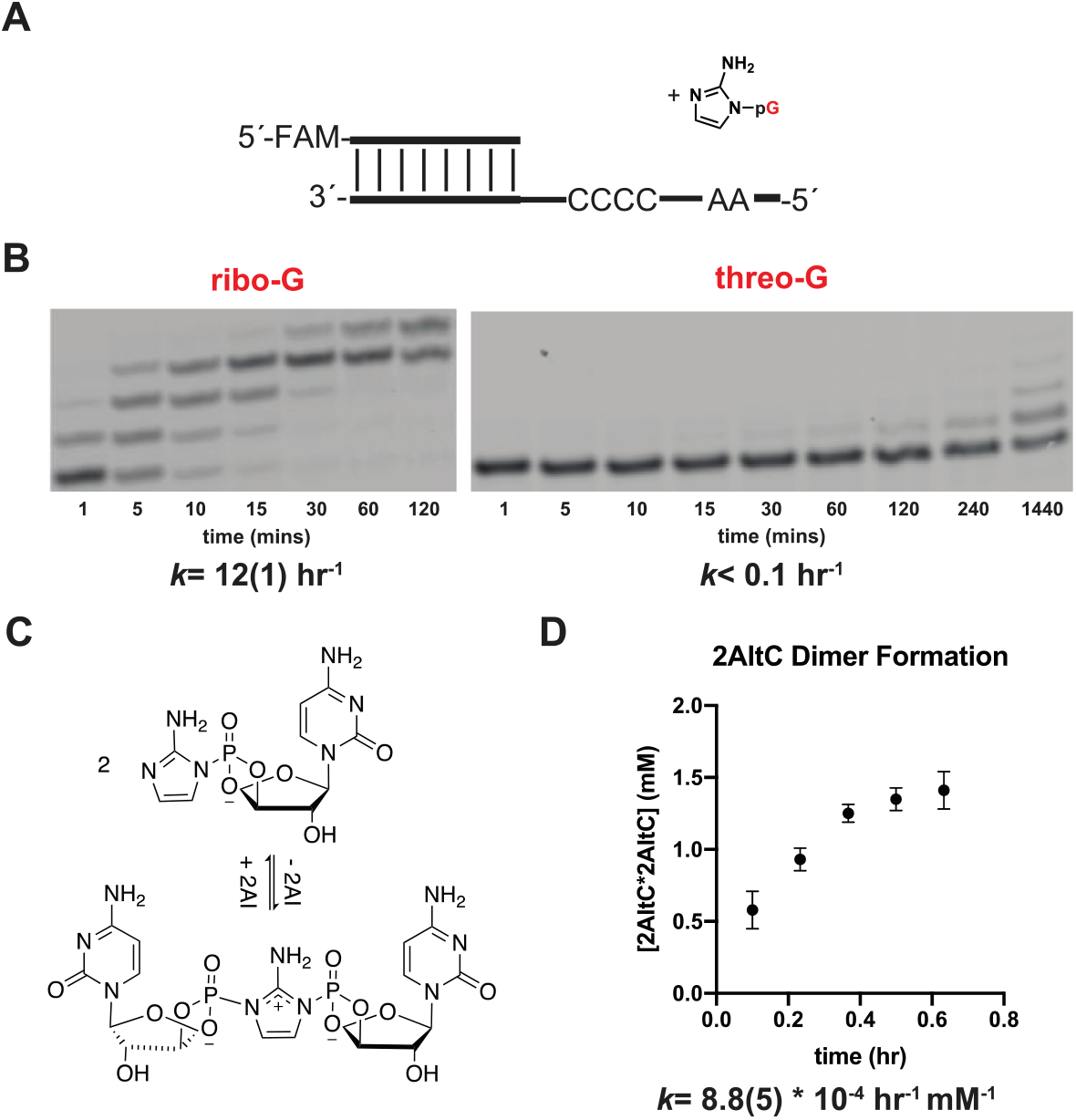
Comparison of nonenzymatic primer extension using either an activated ribonucleotide or an activated threo-nucleotide. (A) Schematic representation of a primer extension reaction with either an activated ribo- or threo-nucleotide. (B) Gel electrophoresis images and rates of primer extension for either an activated ribo-nucleotide (left) or an activated threo-nucleotide (right). All reactions were performed at pH 8.0, in 200 mM HEPES, 200 mM Mg^2+^, 40 mM 2AIpG. Values are the mean rate constant ± SD in parentheses, with the last digit reported being the last significant figure and the one in which error arises from triplicate experiments. (C) Schematic representation of the dimerization equilibrium of 2-aminoimidazole activated threo-C. (D) Accumulation of imidazolium-bridged threo-C at 60 mM 2AItC over the first 38 minutes. The observed initial rate of the formation of imidazolium-bridged threo-C is 8.8*10^-4^ hr^-1^ mM^-1^. See Supplementary Data for kinetics analysis of primer extension, dimer formation, and hydrolysis (Supplementary Data, Figure S-4-5, 11-14).

We have previously shown that nonenzymatic RNA primer extension is mediated by a highly reactive imidazolium-bridged dinucleotide intermediate(25). We therefore attempted to observe the formation of the corresponding TNA dinucleotide intermediate under primer extension conditions (Figure 2C). When we incubated purified 2AIptC at a concentration of 60 mM at pH 8 and followed the reaction by 31P-NMR, we found that the imidazolium-bridged TNA dinucleotide is formed, but the observed rate of 8.8 * 10^-4^ hr^-1^ mM^-1^ is approximately 4 to 5 fold slower than the corresponding rates for the activated ribonucleotides 2AI-rA and 2AI-rC (3.2 and 4.5 * 10^-3^ hr^-1^ mM^-1^)(26,27) (Figure 2D). This result suggests that the slow rate of primer extension with activated threo-nucleotide monomers is due at least in part to the slower formation of the necessary imidazolium-bridged intermediate.

#### 2-AIptG forms an imidazolium-bridged intermediate that binds to the template

Nonenzymatic template-directed primer extension with activated ribonucleotides proceeds through a covalent imidazolium-bridged dinucleotide intermediate(25). As shown above, activated tG monomers do form an equivalent intermediate, albeit more slowly. In order to visualize the structure of the TNA intermediate, and to gain insight into the reasons for its slow formation and possibly slow reactivity with the primer/template complex, we undertook a crystallographic analysis. In prior work, we obtained direct structural evidence for the imidazolium-bridged RNA intermediate (Gp-AI-pG) by first co-crystallizing an RNA-dGMP complex and then exchanging the unactivated nucleotides with an activated guanosine-5’-phosphoro-(2-aminoimidazolide) by soaking the crystals in a solution of activated monomer(28). Here we carried out a similar procedure by soaking the same RNA-dGMP co-crystals with a solution of the activated threo-nucleotide monomer 2-AIptG. Using a self-complementary LNA-modified RNA sequence, 5’-*mCmCmCG*ACUUAAGUCG-3’ (Figure 3A), we sought to observe formation of the dinucleotide intermediate on the 5’-*mCmC* binding site on the template. We screened a range of soak times to estimate the appropriate timescale for exchange and intermediate formation. The optimal structure was determined after soaking the RNA-dGMP crystal in a 20 mM 2-AIptG solution for 24 h. At each end of the A-helical duplex, we observe density consistent with imidazolium-bridged TNA guanosine dinucleotides (tGp-AI-ptG) bound to the template (TNA-D1, Figure 3B and 3C). It is noteworthy that formation of the tGp-AI-ptG intermediate in the crystal was much slower than formation of the corresponding RNA intermediate Gp-AI-pG (24 h vs ~1 h)(28), consistent with our solution phase NMR observations. Both guanine nucleobases in the imidazolium-bridged TNA intermediate tGp-AI-ptG are well ordered and are bound to the template through Watson-Crick base pairing. Critically, the electron density between the two nucleotides is consistent with the formation of an imidazolium bridge. The threose sugars are only moderately well-ordered in the complex, making it difficult to draw conclusions about the sugar conformation in Gp-AI-pG. Nevertheless, the phospho-imidazolium-phospho bridge is sufficiently well ordered to enable an estimate of the attack geometry. The distance between the 3’-OH of the primer and the incoming P atom of tGp-AI-ptG molecule is ~ 5.9 Å in the refined model. This distance is significantly longer than that observed in the RNA complex containing native Gp-AI-pG (4.6 Å)(28). Furthermore the O-P-N angle of attack is 122°, which is less than the previously observed angles of 132° and 170° for the all RNA complex. Both the longer distance and smaller angle could contribute to the slower primer extension observed with activated threo-nucleotides.

**Figure 3.**
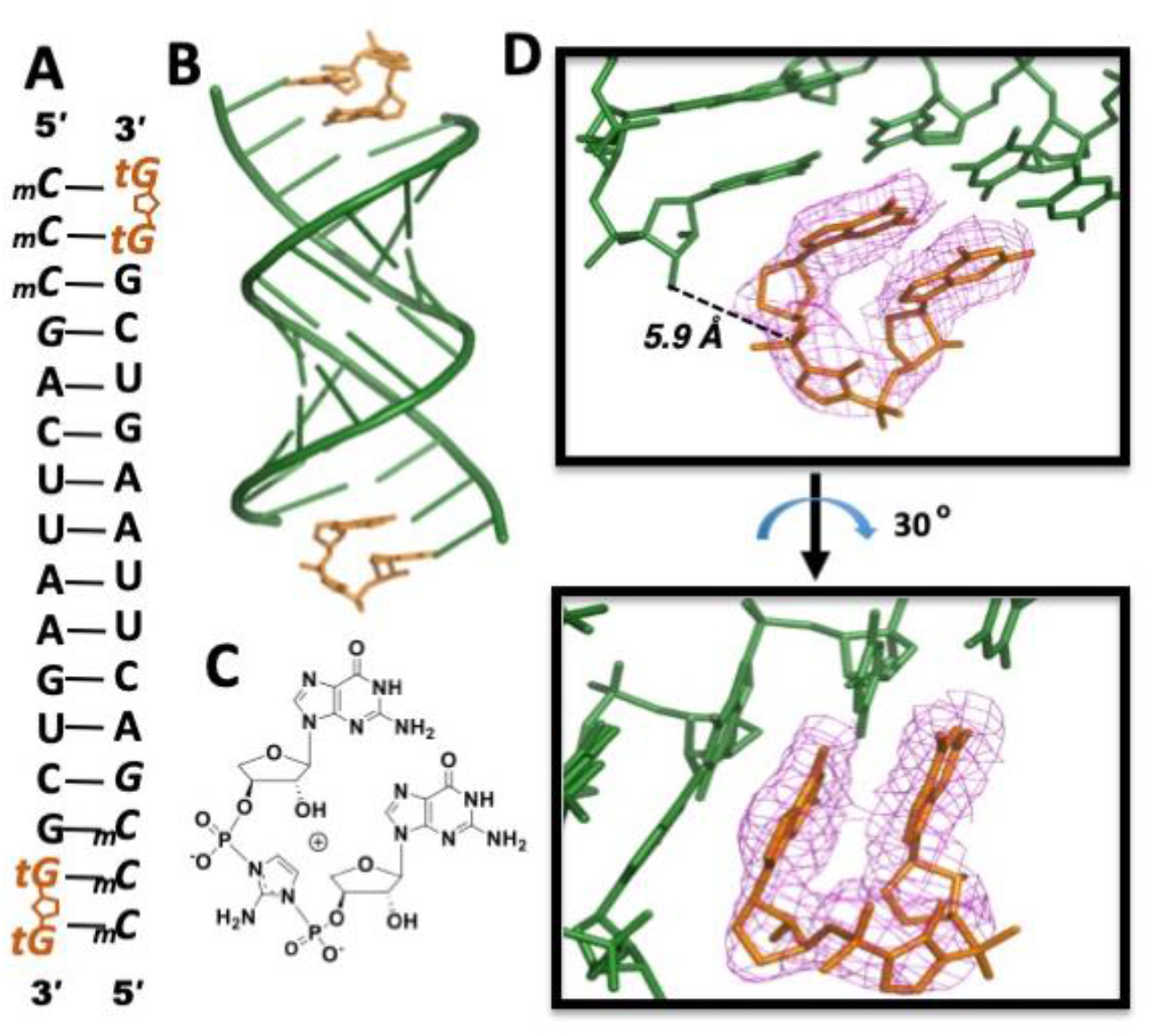
Structure of the imidazolium-bridged tG-dinucleotide intermediate bound to an RNA primer/template complex. (A) Diagram and designed secondary structure of RNA primer/template complex. The imidazolium-bridged tG-dinucleotide intermediate in primer extension is shown bound to the template overhang. The tG 3’-p2AIp-3’ dinucleotides (orange) are bound at both ends of the duplex. (B) Overall crystal structure of TNA-D1 complex. (C) Chemical structure of the imidazolium-bridged threo-guanosine dinucleotide intermediate. (D) Local views of the tG-intermediate bound to the template by two Watson-Crick base-pairs. Pink mesh indicates the corresponding 2*F_o_-F_c_* maps contoured at 1.5 σ. The distance between the 3’-OH group of the primer and the P atom of the adjacent tG of the bound intermediate is 5.9 Å, and the O-P-N angle of attack is 122°.

#### Primer extension with 2-AIptG and an activated downstream helper oligoribonucleotide

Although primer extension with multiple sequential TNA monomers is extremely slow, we considered the possibility that an isolated TNA monomer adjacent to an activated RNA oligonucleotide might show improved reactivity. For example, the formation of an imidazolium-bridged TNA-RNA intermediate might be faster if the imidazole moiety of 2AIptG could attack the phosphate of an adjacent 2AIpG more readily due to the reduced steric bulk around the ribonucleotide phosphate, which is held away from the sugar by C5’. We examined the kinetics of primer extension in this format using an RNA primer/template duplex, with binding sites on the template for 2AIptG and an adjacent 2AI activated RNA trimer (Figure 4). In this context, the rate of primer extension with the activated threo-nucleotide 2AIptG was only four-fold slower than with the corresponding ribonucleotide 2AIpG (Figure 4 B). This simplest explanation for this result is that mixed TNA-RNA intermediates both form and react more rapidly than TNA-TNA intermediates, permitting threo-nucleotide incorporation with only a modest kinetic defect.

**Figure 4.**
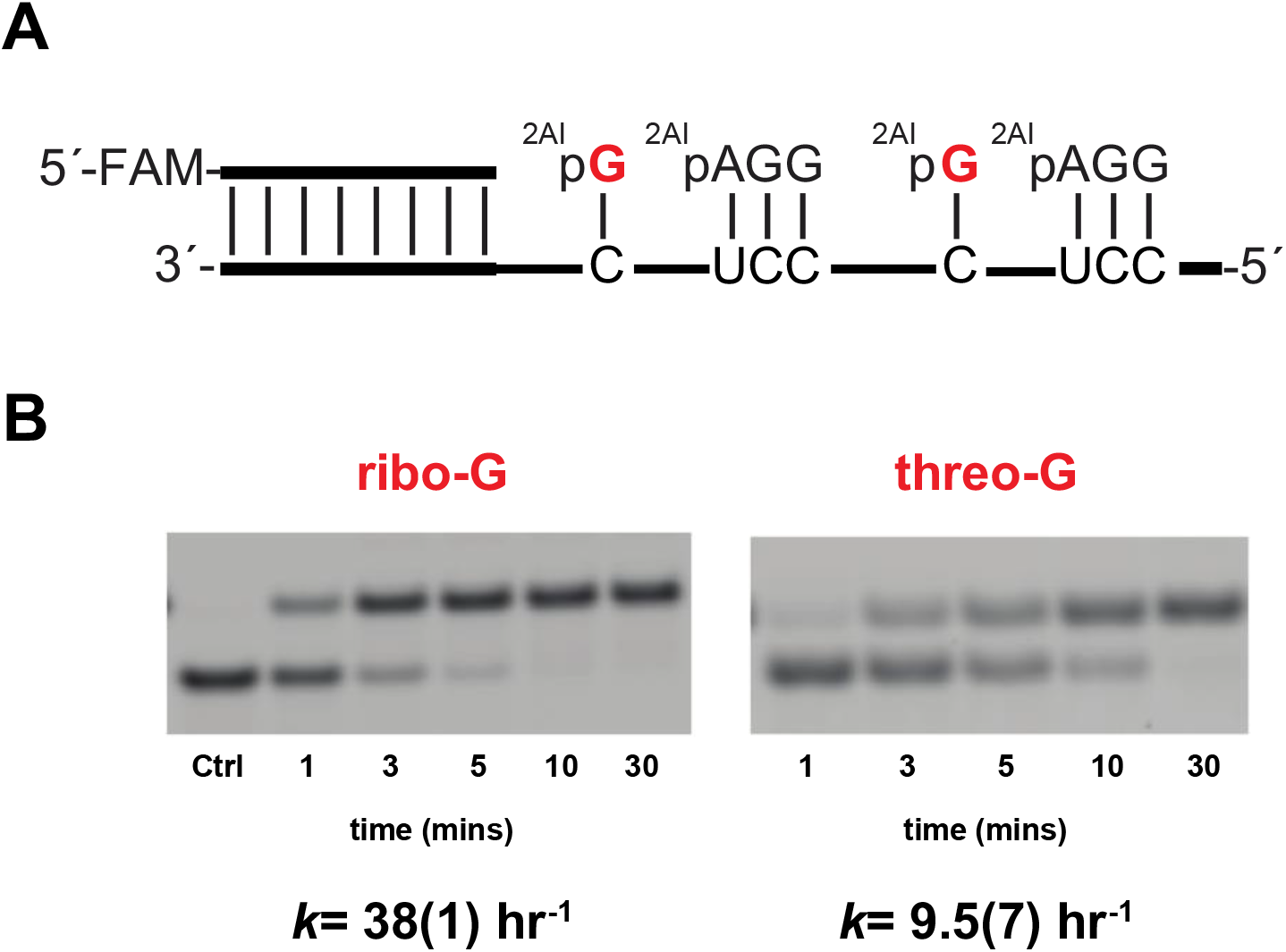
Evaluation of threose nucleotides in nonenzymatic primer extension. (a) Schematic representation of primer extension using an RNA template. 2AIpG represents 2-aminoimidazole activated guanosine monomer and 2AIpAGG represents 2-aminoimidazole activated RNA trimer helper. (b) Gel electrophoresis images and rates of primer extension for 2-aminoimidazole activated nucleotides. All reactions were performed at pH 8.0, 200 mM HEPES, 200 mM Mg^2+^, 20 mM 2AIpG, 1 mM of activated RNA trimer. Values are reported as the mean rate constant ± SD in parentheses, with the last digit reported being the last significant figure and the one in which error arises from triplicate experiments. See Supplementary Data for kinetics analysis of primer extension (Supplementary Data, Figure S-6 and 7).

To further investigate possible reasons for poor primer extension with threo-nucleotide monomers, we obtained two additional structures of RNA oligonucleotides with bound threo-nucleotide monomers. We first co-crystallized the unactivated α-L-threofuranosylguanine-3’-monophosphate (tGMP) monomer with a short partially self-complementary RNA 7-mer (TNA-M1, sequence: 5’-*mCmUG*UACA-3’, italic: locked nucleotides, *mC:* locked 5-methylcytidine, *mU:* locked 5-methyluridine, Figure S-2. This oligonucleotide self-anneals to form a central six base pair duplex, flanked by single overhanging 5-Me-C residues, to which the complementary G or tG monomer can bind. When incubated with tGMP, crystals grew in the hexagonal P6_3_ space group, and we determined the structure to a resolution of 2.36 Å. Similar to our previously reported RNA-GMP complex structure(29), the helical duplexes are end-to-end stacked, and a tGMP nucleotide is bound to the templating 5-methylcytidine residue at each end in a Watson-Crick base pair (Figure S-2B). The sugar is in the C4’-*exo* conformation in both tGMP ligands (Figure S-2C). The well-defined electron density of the 3’-phosphate of tGMP shows that the distance between the 3’-OH of the RNA primer and the phosphorus atom of tGMP is about 6.7 Å, slightly longer than the distance observed for the ribo-nucleotide monomer (~6.3 Å)(29).

To examine whether a structured downstream oligonucleotide might help to preorganize the noncovalently bound tGMP, an effect we have previously observed with RNA monomers, we co-crystallized a tGMP monomer sandwiched within a gapped RNA hairpin structure(30,31). The resulting structure (TNA-M2, Figure S-2D and E) reveals that the sandwiched tGMP nucleotide is in a well-ordered Watson-Crick pair, and the tGMP sugar is in the C4’-*exo* conformation (Figure S-2F). Based on the well-defined electron density of the tGMP 3’-phosphate, the distance between the 3’-OH of the primer and the incoming phosphorus atom is ~4.6 Å, similar to but slightly longer than the corresponding distance observed for a bound ribonucleotide (~4.4 Å) (30). In both structures, the longer distance between the primer 3’-OH and the bound tGMP monomer appears to be influenced by the structure of the threo-nucleotide, in which the phosphate group is directly attached to the threo-furanosyl ring instead of being held away from the sugar by the 5’-methylene of ribose.

One noteworthy conformational difference between complexes TNA-M1 and TNA-M2 is that the bound monomers have χ torsion angles that differ by >40°. Without any helper to enhance the binding, the χ angle of tGMP in TNA-M1 is −177°, which is closer to the characteristic χ torsion angle in an A-form helix. When the threo-nucleotide is sandwiched between two well-structured RNA duplexes, the N-glycosidic bond of the TNA monomer is rotated, resulting in a χ angle of −134°. This latter angle is closer to those observed for the covalently incorporated TNA residues in the TNA-P1 and TNA-T1 (see below) stem-loop structures (Table 1), suggesting that the conformational constraints imposed by the sandwiched RNA complex induce a threo-nucleotide geometry that is distinct from that of the free species. Moreover, the threo-nucleotide monomers in TNA-M1 and TNA-M2 have different γ torsion angles from the backbone-linked threo-nucleotides in TNA-P1 and TNA-T1 (Table 1). In the absence of covalent linkage to the RNA backbone at either the 3’ or 2’ terminus, the tGMP monomers in both TNA-M1 and TNA-M2 complexes may be in a more relaxed conformation, to differing extents depending on the presence of a proximal downstream oligonucleotide.

**Table 1.** Torsion angles in degrees of TNA residues. (na = not available)

| Complex | Status of Residue | $\alpha$ | $\gamma$ | $\delta$ | $\epsilon$ | $\zeta$ | $\chi$ |
| --- | --- | --- | --- | --- | --- | --- | --- |
| TNA-P1 | primer | -76.5 | -177.4 | 163.6 | na | na | -121.3 |
| TNA-T1 | template | -75.4 | -177.1 | 140.1 | -96.2 | -111.4 | -111.4 |
| TNA-M1 | paired monomer | na | -137.1 | 172.0 | na | na | -177.2 |
| TNA-M2 | paired monomer | na | -138.2 | 171.4 | na | na | -134.7 |
| 2'-NH<sub>2</sub>tG(13) | free monomer | na | na | 163.1 | na | na | -138.2 |

### Kinetic and structural consequences of TNA residues in the primer and the template

#### Effect of a threo-nucleotide at the primer terminus

We then asked whether nonenzymatic extension beyond a terminal threo-nucleotide at the end of an RNA primer was observable. Our previous results with 2’-NH_2_ modified threo-nucleotides suggested that primer extension from a threo-nucleotide should be possible (14). However, because the amino sugar modification is known to affect nucleotide conformation(32) and because the amino group is more nucleophilic than a hydroxyl (33), it was unclear how well primer extension would proceed from an unmodified threo-nucleotide. We therefore synthesized an RNA primer containing a single terminal threo-cytidine (tC) residue at its 3’ end (2’ end when tC is in the terminal position, 5’-(FAM)-AGUGAGUAACGtC-2’, Figure 5A). We then compared rates of RNA-templated primer extension using activated 2-AIpG monomer as a substrate for the extension of this tC-terminal primer vs. an all-RNA control primer. In the control RNA primer/template system, primer extension proceeded rapidly, with an observed rate of 12 h^-1^. With the tC-terminal primer, however, the observed rate was reduced to 1.2 h^-1^, 10-fold slower than the RNA extension reaction. After incubation for 24 hours, the tC-terminated primer was primarily converted a +4 nucleotide product (Figure 5B). Thus, once the primer is extended with a ribonucleotide, continued primer extension can proceed to the end of the template.

**Figure 5.**
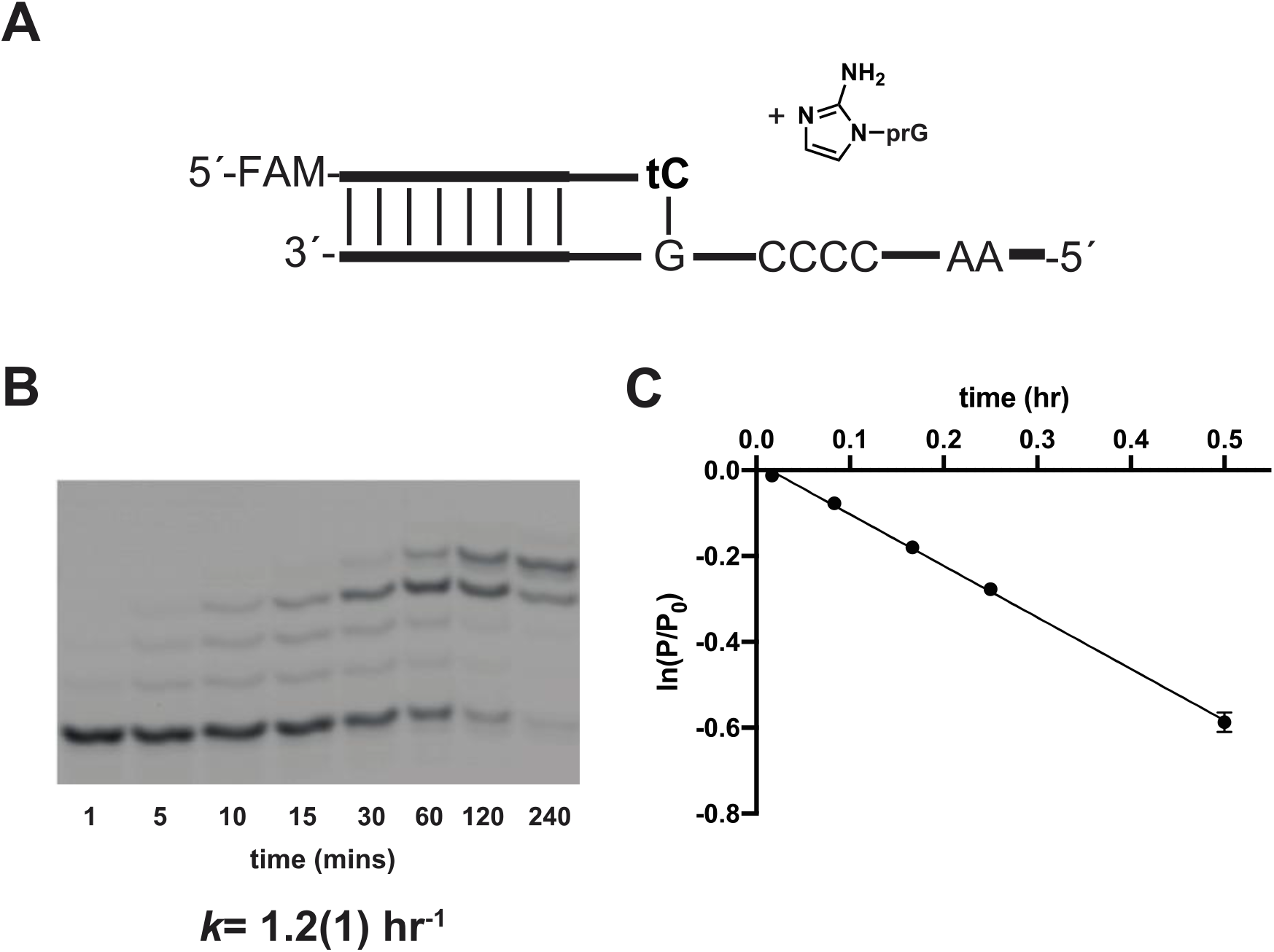
Primer extension with an RNA primer ending in a threo-nucleotide. (a) Schematic representation of a primer extension reaction. Primer extension reactions were carried out in triplicate using 40 mM 2AIprG, pH 8.0, in 200 mM HEPES, 200 mM Mg^2+^. (b) Representative PAGE analysis of result. Values are the mean rate constant ± SD in parentheses, with the last digit reported being the last significant figure and the one in which error arises from triplicate experiments. (c) Plot of ln(P/P_0_) as a function of time. The rate of extension was determined from linear least-square fits of the data from three independent experiments.

#### Structural consequences of a single threo-nucleotide at the primer terminus

To gain insight into the structure of a reaction center containing a priming threo-nucleotide, we crystallized an RNA hairpin-monomer complex (TNA-P1, Figure 6A), in which the RNA stem contained a 2’ terminal threo-guanosine (tG) residue followed by a single-nucleotide gap. One unactivated cytidine monophosphate (CMP) mononucleotide was sandwiched noncovalently between the tG primer and the downstream helper oligonucleotide, to mimic the bound monomer in the ground state prior to primer extension. As in our previously determined RNA-monomer hairpin structure(34), the entire RNA-monomer hairpin crystallized in the R3 space group, with one complex per asymmetric unit (Figure 6B). We determined the structure to a resolution of 2.7 Å. The tG-containing RNA, downstream RNA helper and bound CMP monomer assembled to form the desired hairpin, with the tetraloop of one hairpin docked with the receptor motif of a separate hairpin complex. The terminal tG residue at the 2’-end of the primer was well ordered, with the guanine nucleobase Watson-Crick base-paired with the templating cytidine. The threose sugar is in a C4’-*exo* conformation with the 2’ and 3’ hydroxyls pseudo-diaxial (Figure 6C), consistent with a prior report from Egli et al. in which a TNA residue was present in the middle of an oligonucleotide(8). The 2’-OH group of the tG residue is oriented toward the phosphate of the incoming CMP monomer, which is Watson-Crick paired with the template. The distance between the P atom of the bound CMP and the 2’-OH of the tG primer is ~ 3.2 Å, which appears favourable for attack, however as the monomer is unactivated CMP, the structure is not informative about the angle of attack with respect to the leaving group.

**Figure 6.**
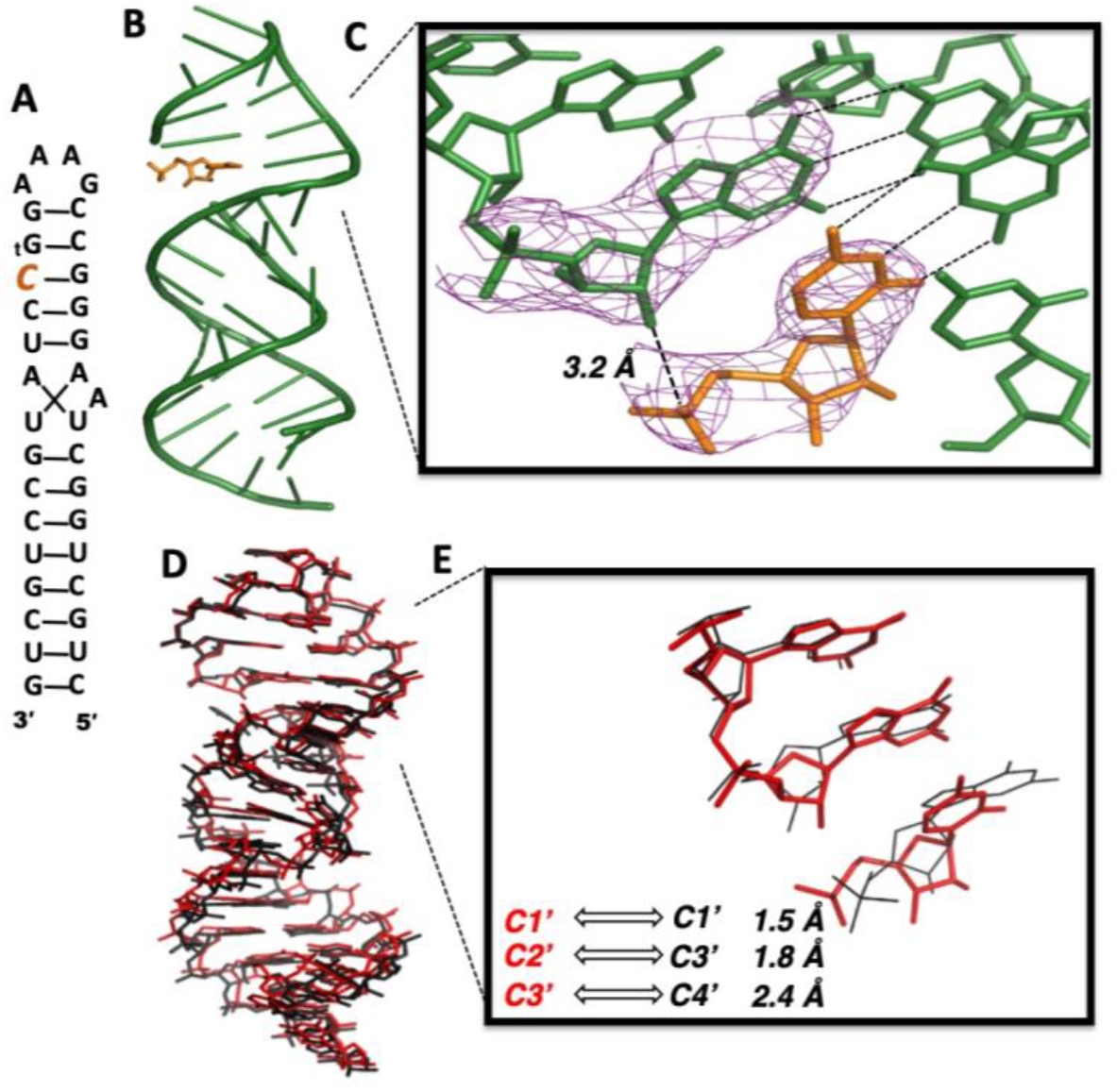
Threo-nucleotide substitution at the end of an RNA primer has minor structural effects. (A) Diagram and designed secondary structure of RNA hairpin-monomer complex TNA-P1. The primer and template are linked together to form a hairpin structure. A CMP monomer (orange) is sandwiched between the TNA guanosine (tG) at the end of the primer and the flanking downstream oligonucleotide. (B): Overall crystal structure of TNA-P1 complex. Green: primer-template hairpin and downstream RNA oligonucleotides. Orange: CMP monomer. (C) Local view of the terminal primer tG residue and the adjacent CMP monomer. Pink mesh indicates 2*F_o_-F_c_* maps contoured at 2.5 σ. (D) Side view of the superimposed structures of the previously reported all RNA hairpin structure (5VCI) and the TNA-P1 complex. Red: TNA-P1 structure; black: 5VCI structure. (E) Local view of the superimposed structures. Distances between corresponding atoms in the threose sugar and ribose sugar versions are labelled (middle nucleotide of overlaid structures).

The overall structure of the TNA-P1 complex is highly similar to that of the RNA hairpin-monomer complex presented in our recent report (PDB 5VCI)(34). The two structures are fully superimposable (RMSD=0.96 Å), including the RNA backbone and nucleobase positions (Figure 6D). Thus, a single threo-nucleotide at the end of an RNA duplex is accommodated with minimal structural perturbation. The largest deviations between the two structures are at the primer terminus, where the tG residue is shifted and twisted compared to the native RNA primer, with distances in the range of 1.5 - 2.4 Å between the corresponding atoms of the threose and ribose sugar rings (Figure 6E). The tG residue has a glycosidic linkage torsion χ angle of −121° (Table 1), compared to a typical χ angle of −168° for the ribonucleotides of the primer(26).

#### Primer extension on templates containing TNA nucleotides

We next sought to investigate the effect on primer extension of single or multiple TNA residues in the template. Previous work from Switzer et al. showed that primer extension across an all TNA template region was possible but slow(12). We therefore began by comparing primer extension on a template region consisting of four ribo- or threo-cytidine residues, using an all RNA primer and activated ribo-guanosine (2AI-pG) as the monomer substrate (Figure S-3). Consistent with the previous results of Switzer et al. (13), primer extension on the (tC)_4_ template (1.3 h^-1^) was roughly ten-fold slower than on a (rC)_4_ template (12 h^-1^). (Figures S-3B and S-10)

In a prebiotic situation in which both ribo- and threo-nucleotides were present, templates in which isolated threo-nucleotides were interspersed in an otherwise RNA oligonucleotide might be common. However, the effect of an isolated threo-nucleotide in an otherwise all-RNA template has not been previously investigated. We therefore synthesized two new template oligonucleotides (5’-AAGGG(t)GCCAGUCAGUCUACGC-3’), one all RNA and one containing a single threo-guanosine nucleotide (tG) at the +1 position (Figure 7). When the tG residue was present in the template strand at the +1 position the pseudo-first order reaction rate for the first addition was 5.6 h^-1^, compared to 11 h^-1^ for the all-RNA template (Figure 7B). However, extension of the +1 product to longer products was much slower (estimated rate constant for conversion of +1 to +2/3/4 products 0.71 h^-1^), a decrease comparable to that see for a TNA residue at the end of the primer. Thus, a TNA:RNA base-pair seems to significantly slow primer extension, irrespective of whether the TNA residue is in the primer or the template. However, subsequent extension to the +3 and +4 products was not observably inhibited, and almost complete conversion of the primer to fully extended +4 products was observed after 24 hours.

**Figure 7.**
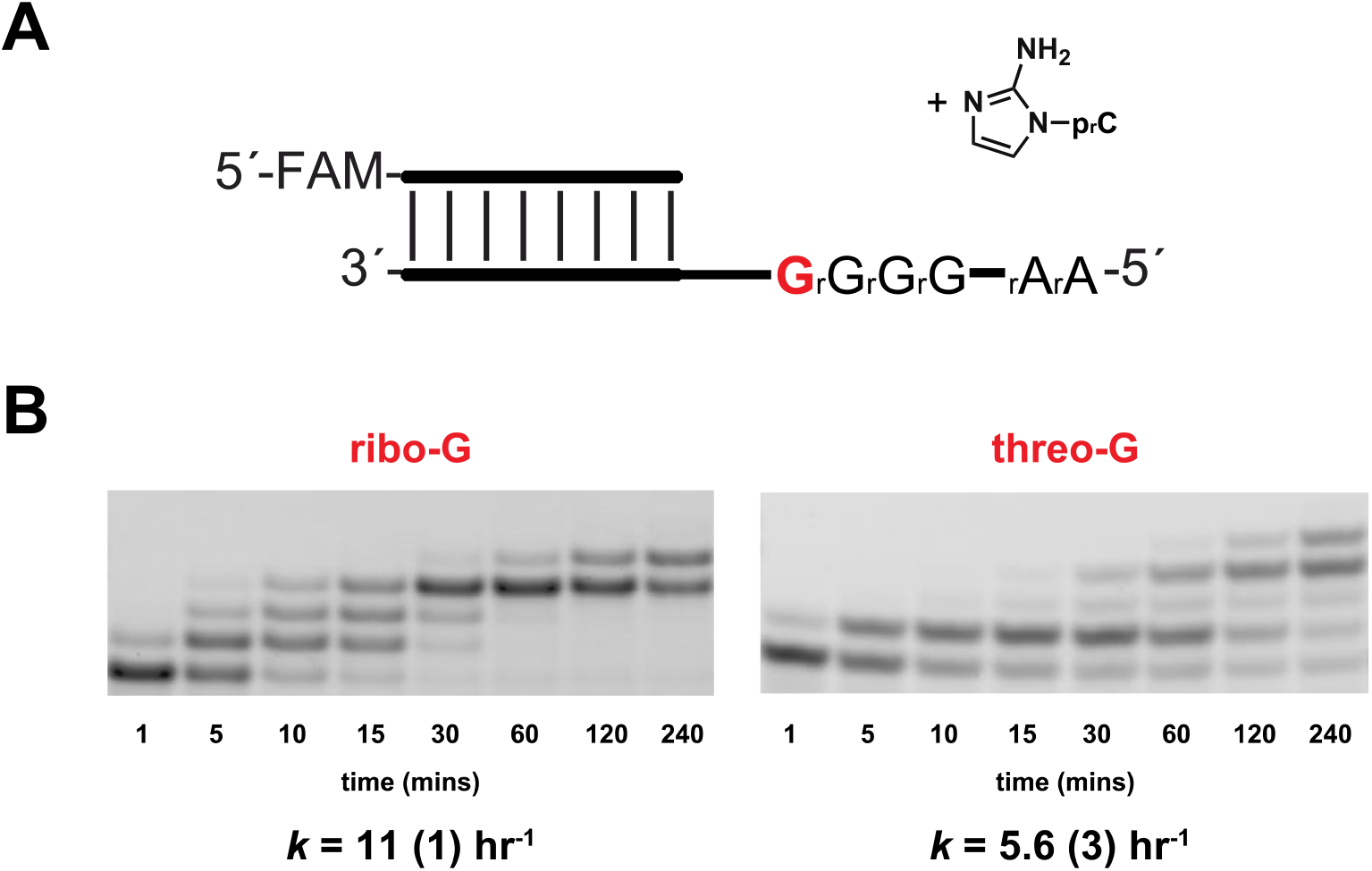
Evaluation of nonenzymatic primer extension with 2-aminoimidazole activated ribo-cytidine (2AIpC) across from a single templating threo-guanosine (tG) residue. (a) Schematic representation of a primer extension reaction of 2AIpC with template containing either ribo-G (rG) or threo-G (tG). (b) Gel electrophoresis images and rates of primer extension with 2-aminoimidazole activated cytidine. All reactions were performed at pH 8.0, 200 mM HEPES, 200 mM Mg^2+^, 40 mM 2AIpC. Values are reported as the mean rate constant ± SD in parentheses, with the last digit reported being the last significant figure and the one in which error arises from triplicate experiments. See Supplementary Data for kinetics analysis of primer extension (Supplementary Data, Figure S-8 and 9).

#### Structural consequences of a single TNA residue in the template strand

To gain structural insight into the conformation of the reaction center in TNA-templated nonenzymatic primer extension, we co-crystallized and determined the structure of an RNA hairpin complex, similar to that used above, but this time containing a single TNA residue on the template strand of a gapped stem. The RNA hairpin-monomer complex (TNA-T1) contains a single tG residue at the appropriate position to serve as the template to bind a free CMP monomer (Figure 8A). Broadly, TNA-T1 and TNA-P1 shared similar structural features, including an RNA A-form stem, tetraloop/receptor motif, and sandwiched monomers (Figure 8B). The TNA-T1 complex also crystallized in the R3 space group, and we determined the structure to a resolution of 2.95 Å. The tG residue on the template strand is well-ordered, forming three hydrogen bonds with the bound CMP monomer through a Watson-Crick base-pairing interaction. The tG threose sugar ring is in the C4’-*exo* conformation, and the glycosidic torsion angle x of tG is −111°, similar to the tA residue seen at the end of the primer. However, the noncovalently bound CMP is poorly ordered, suggesting a significant degree of conformational heterogeneity. The electron density of the monomer’s sugar and phosphate is poorly resolved, yielding only a rough estimate of 4 Å for the distance between the 3’-OH of primer and the P atom of the CMP (Figure 8C).

**Figure 8.**
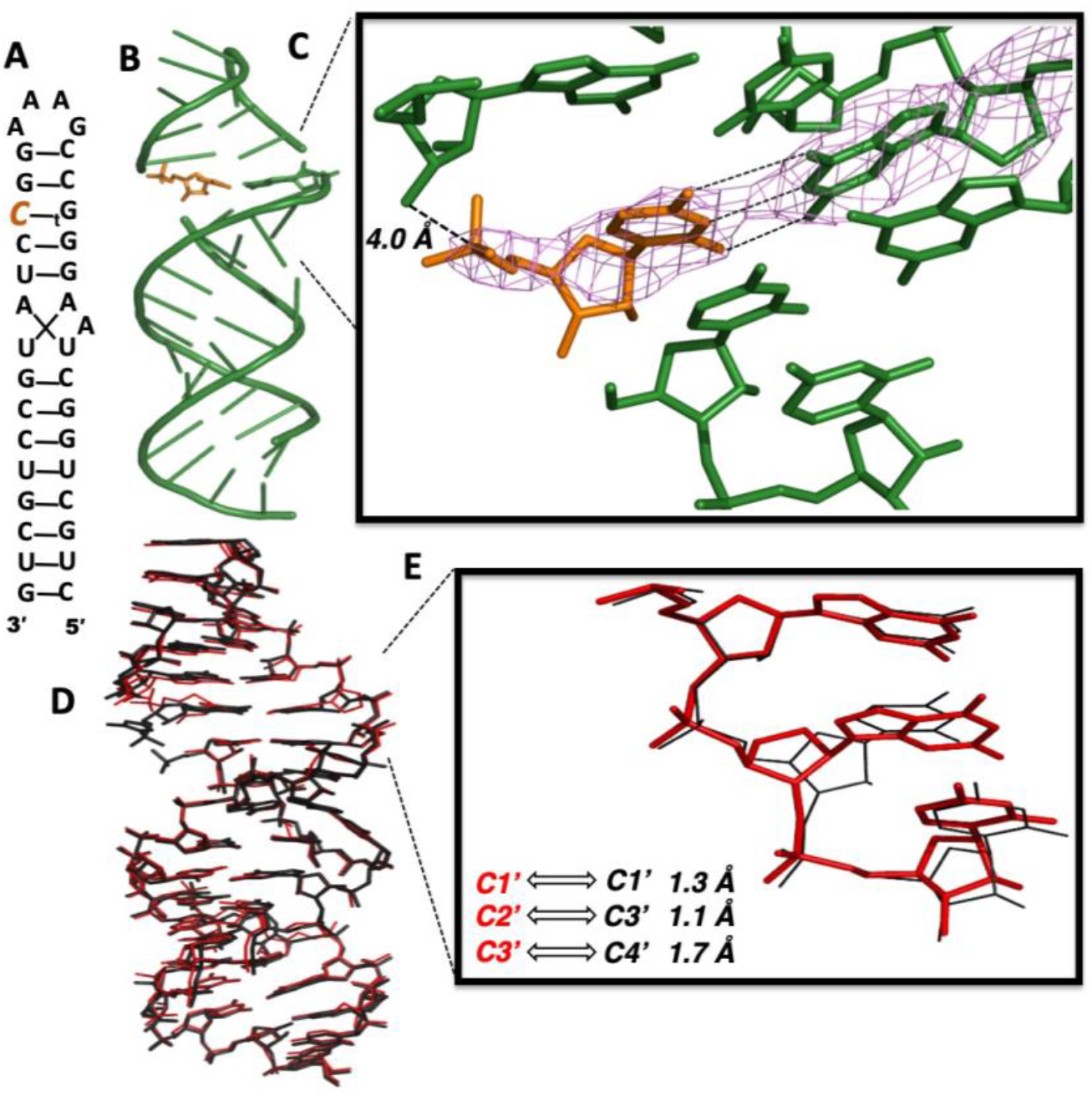
A threo-nucleotide substitution in an RNA template results in only minor and local structural changes. (A) Diagram and designed secondary structure of RNA hairpin-monomer complex. A CMP monomer (orange) is sandwiched between the 3’-end of the primer and a downstream oligonucleotide. The TNA substitution in the template is denoted by tG (B): Overall crystal structure of TNA-T1 complex. (C) Local view of the tG template residue, showing Watson-Crick base-pairing to the bound CMP monomer. Pink mesh indicates the corresponding 2*F_o_-F_c_* maps contoured at 2.5 σ. (D) Side view of the superimposed structures of the previously reported RNA hairpin structure (5VCI) and the TNA-T1 complex. Red: TNA-T1 structure; black: 5VCI structure. (E) Local view of the superimposed structures. The distances between the positions of the corresponding atoms in the threose sugar and the corresponding ribose sugar are labelled.

Superimposing the TNA-T1 structure over the all-RNA complex 5VCI, we find the two complexes are very similar in overall conformation (RMSD = 0.64 Å, Figure 8D). The tG threose sugar, constrained by stacked upstream and downstream RNA nucleotides, is shifted by 1.1-1.7 Å compared to the native RNA ribose sugar ring (Figure 8E). The torsion angles of the tG backbone are presented in Table 1. As expected, the tG residues in the TNA-P1 and TNA-T1 complexes share similar α and γ torsion angles, likely due to similar structural constraints on the threo-nucleotide residue in the RNA context of both hairpin structures. Overall, a single TNA nucleotide substitution on the template does not dramatically disrupt the RNA structure.

## DISCUSSION

For many years the RNA World hypothesis has been a guiding force for studying the transition from simple chemistry to complex cellular life, bolstered by recent advances in prebiotically plausible pathways for the generation of RNA building blocks from simple feedstocks(35,36) and in the nonenzymatic copying of RNA templates in prebiotically plausible compartments(37). At the same time, we and others have explored alternative genetic materials with the potential to store and replicate genetic information(1). Among these alternatives, TNA is a striking candidate that, because of its simpler chemistry, might present a lower hurdle for abiogenesis(4). However, TNA appears to have poor reactivity in nonenzymatic copying reactions, regardless of whether the threo-nucleotide 2’-position is the native-OH group, as shown in our current work, or the highly nucleophilic-NH_2_ group(14).

Our structural studies provide reasonable explanations for the moderately reduced rates of primer extension from a primer ending in a threo-nucleotide, and across from a threo-nucleotide in the template strand. In the corresponding interrupted RNA hairpin complexes TNA-P1 and TNA-T1, which have single threo-nucleotide substitutions at the end of the primer or in the template strand, the overall conformations are generally comparable to those of the all-RNA complexes. In this stem-loop RNA context, TNA residues adopt an extended conformation so as to be accommodated within the RNA A-form duplex with minimal global distortion, and with the formation of well-ordered base pairs with complementary RNA nucleotides on the opposite strand. These structural findings are in agreement with the observation that TNA strands form thermodynamically stable heteroduplexes with RNA(4,8). The modeled 2’-O to 5’-P distance for nucleophilic attack in the case of the threo-nucleotide terminated primer is also comparable to that seen in fully RNA complexes. Given that duplex structure is unperturbed and that the local structure is only minimally distorted, it is reasonable that only moderately reduced reactivity is observed for ribonucleotide addition to a threo-nucleotide terminated primer or across from a threo-nucleotide in the template.

Remarkably, the extremely slow rate of nonenzymatic primer extension with 2AI-activated threo-nucleotide monomers alone appears to result largely from the failure of activated threo-nucleotides to form significant levels of the imidazolium-bridged dinucleotide intermediate in solution, with a possible further contribution from slower reaction of intermediate that does form with the primer. The reaction of two 2AI-activated threo-nucleotides with each other to form the imidazolium-bridged dinucleotide intermediate involves the attack of the unprotonated N3 of the 2AI moiety of one threo-nucleotide on the phosphate of a second activated monomer, with its protonated 2AI being the leaving group. The low level of imidazolium-bridged dinucleotide seen in solution was at first surprising, because the reacting groups are distant from the threose sugar that distinguishes TNA from RNA. However, the phosphate of an activated threo-nucleotide is more sterically hindered than that of a ribo-nucleotide, because the TNA phosphate is directly linked to the threo-furanosyl ring such that it is flanked by the nucleobase and the sugar as well as the covalently attached 2AI moiety. We speculate that this steric hindrance is responsible for the low levels of imidazolium-bridged TNA intermediate in solution, and hence the slow rate of primer extension with 2AI-activated threo-nucleotide monomers. Nevertheless our observations show that the imidazolium-bridged TNA intermediate can form *in crystallo*, similarly to activated RNA monomers(28), albeit much more slowly. The formation of the imidazolium-bridged TNA dinucleotide in the crystal may be facilitated when two activated threo-nucleotide monomers are held in close proximity for extended periods of time as a result of being bound next to each other on the template. Additional experiments will be required to identify all of the factors that control the formation of the imidazolium-bridged TNA intermediate both in solution and *in crystallo*.

Our crystallographic findings show that both TNA monomers and the TNA imidazolium-bridged dinucleotide do bind to the template by Watson-Crick base pairing, and the threose sugar is consistently in the C4’-*exo* conformation. The unactivated 3’-phosphate in single threo-nucleotides is shifted towards the helix major groove, increasing the distance from the attacking 3’-OH of the primer. Compared to single TNA residues in the primer or template, the γ torsion angles of free monomers adopt distinct values that influence the orientation of the 3’-phosphate group. The noncovalently bound TNA monomer, unconstrained by linkage to the RNA backbone, appears to relax to a conformation that is unfavorable to primer extension. In the case of the imidazolium-bridged dinucleotide, the threose sugars are disordered, but the appearance of electron density between the two threo-nucleotides clearly indicates the formation of the critically important imidazolium-bridge. The estimated distance between the primer 3’-OH and the P atom of the adjacent TNA nucleotide is significantly longer than for a ribonucleotide (5.9 Å vs 4.6 Å), suggesting that even if 2AI-activated TNA monomers do form the intermediate and bind the template, primer extension would be very slow.

Our results cast doubt upon the viability of TNA as a self-replicating polymer, at least in the context nonenzymatic template-directed primer extension with phosphorimidazolide-activated threo-nucleotide monomers. The copying of a C4 template region with 2AI activated tG monomers was extremely inefficient no matter whether the primer ended in a ribo- or a threo-nucleotide, and no matter whether the template region was composed of a ribo- or a threo-nucleotides. Since even copying in an essentially all TNA context was ineffective, the inability of 2AIptG monomers to mediate efficient multi-step primer extension cannot simply reflect the local structural context of the reaction. Instead, the more likely explanation follows from our observation that formation of an imidazolium-bridged tG-p2AIp-tG dinucleotide is inefficient, such that the low levels of this key intermediate lead to poor primer extension irrespective of the local structural context.

We have previously considered a scenario in which ribonucleotides were the predominant but not the only prebiotic synthetic product, such that heterogeneous oligonucleotides might have assembled by non-templated chemistry. Just as in our previous consideration of arabino-nucleotides, we now find that isolated threo-nucleotides in the template can be copied over by RNA primer extension, but more slowly than an all RNA template. Similarly, a primer ending in a single threo-nucleotide can be extended with ribonucleotide monomers, but with a ten-fold kinetic defect compared with an all RNA primer. Interestingly, the kinetic defect associated with an arabino-terminated primer is more severe (~ 100-fold). In contrast, the kinetic defect associated with primer extension with threo-nucleotide monomers is context dependent. Our results such that the major route for the incorporation of threo-nucleotides into a growing chain would be through reaction of the primer with a mixed TNA-RNA imidazolium-bridged intermediate, for which the kinetic defect is only about 4-fold relative to ribonucleotide incorporation. Nevertheless this suggests that in the presence of, for example, an equimolar mixture of activated threo- and ribonucleotides, RNA synthesis would predominate. RNA has an exceptional double helical structure with the ribose sugars in the 3’-*endo* conformation; as a consequence the RNA primer/template/intermediate complex is preorganized so as to favour rapid primer extension. Our experiments show that in this context, RNA copying has a significant kinetic advantage over alternative nucleic acids including not only TNA, but also ANA and DNA(26). Our experimental results are therefore consistent with a model in which relatively homogeneous RNA oligonucleotides emerged from a much more heterogeneous mixture of activated nucleotides through the self-purifying capacity of nonenzymatic RNA replication.

## ACCESSION NUMBERS

Atomic coordinates and structure factors for the reported crystal structures have been deposited with the Protein Data bank under accession numbers 6U7Y, 6U7Z, 6U89, 6U8F, 6U8U.

## SUPPLEMENTARY DATA

Supplementary Data are available at NAR online.

## ACKNOWLEDGEMENTS

We thank the Szostak lab for helpful discussions, insightful commentary and careful revision of the manuscript. We thank Dr. L. Zhou for assistance with LC-MS analysis. X-ray diffraction data were collected at the Advanced Light Source (ALS) SIBYLS beamlines 821 and 822, a national user facility operated by Lawrence Berkeley National Laboratory on behalf of the Department of Energy, Office of Basic Energy Sciences, through the Integrated Diffraction Analysis Technologies (IDAT) program, supported by DOE Office of Biological and Environmental Research. The Berkeley Center for Structural Biology is supported in part by the National Institutes of Health, National Institute of General Medical Sciences, and the Howard Hughes Medical Institute.

