## Supplementary Data for "Structural interpretation of the effects of threo-nucleotides on nonenzymatic template-directed polymerization"

[1] Howard Hughes Medical Institute and Center for Computational and Integrative Biology, Massachusetts General Hospital, Boston, MA 02114, USA
[2] Department of Genetics, Harvard Medical School, Boston, MA 02114, USA
[3] Department of Biochemistry and Molecular Biology, Indiana University School of Medicine, Indianapolis, IN 46202, USA
[4] Department of Chemistry and Chemical Biology, Harvard University, Cambridge, MA 02138, USA
[5] Department of Chemistry and of Pharmaceutical Sciences, University of California, Irvine, CA 92697, USA
[a] Present address: Beam Therapeutics, 26 Landsdowne St., Cambridge, MA 02139, USA

### CONTENTS

1. Materials and Methods S2
2. Supplementary Figures S1-14 S4
3. X-ray Crystallographic Studies S18
4. References S20

### 1. Materials and Methods

**1.1 General Information.** All chemicals were purchased from Sigma-Aldrich (St. Louis, MO) unless otherwise noted. 2-aminoimidazole HCl was purchased from CombiBlocks, Inc. (San Diego, CA). Reverse phase flash chromatography was performed using prepacked RediSep Rf Gold C18Aq 50 g columns from Teledyne Isco (Lincoln, NE).

**1.2 Oligonucleotide Synthesis.** Native, LNA-containing, and TNA-containing RNA oligonucleotides used for primer extension and crystallographic studies were custom-synthesized by Exiqon Inc. (Woburn, MA) or IDT Inc. (San Jose, CA), or were synthesized in-house on an Expedite 8900 DNA/RNA synthesizer. TNA phosphoramidites were synthesized as previously reported(15,16). Oligonucleotides synthesized in-house were deprotected using AMA (1:1 v/v aqueous mixture of 30% w/v ammonium hydroxide and 40% w/v methylamine) for 20 min at 65 °C, followed by desilylation with $Et_3N$•3HF. Oligonucleotides were HPLC purified on an Agilent ZORBAX Eclipse-XDB C18 column using 25 mM triethylammonium bicarbonate in $H_2O$ (pH 7.5) with gradient elution from 0 % to 30 % acetonitrile over 40 mins. The oligonucleotides were collected, lyophilized, desalted, and concentrated as appropriate for primer extension and crystallization experiments. Oligonucleotides were characterized by LC-MS.

**1.3 Determination of RNA concentration.** Concentrations of the aqueous RNA samples were determined by their UV absorption at 260 nm on a Thermo Scientific Nanodrop 2000c Spectrophotometer (Waltham, MA). The theoretical molar extinction coefficients of these samples at 260 nm were provided by Exiqon.

**1.4 Synthesis of Activated Nucleotides (2AIptC and 2AIptG).** 2AIptC and 2AIptG were prepared according to a previously reported procedure(1). (3′-*O*-phosphoro-α-L-threofuranosyl)guanine disodium salt (0.3 mmol, 1 equiv., 100 mg) and 2-aminoimidazole hydrochloride (3 mmol, 10 equiv., 359 mg) were dissolved in 10 mL of deionized water, followed by pH adjustment to pH 6 by concentrated NaOH and lyophilization. In the case of 3′-*O*-phosphoro-α-L-threofuranosyl)cytidie disodium salt, the reaction was performed as described previously without prior lyophilization(1). The lyophilized sample was resuspended in 10 mL DMSO in an oven-dried, argon protected round-bottomed flasks with a pre-dried magnetic stir bar. Triphenylphosphine (3 mmoles, 10 equiv., 787 mg) and anhydrous triethylamine ($\rho$ = 0.726 g $mL^{-1}$, 3 mmoles, 10 equiv., 418 µL) was sequentially charged. 2,2′-dipyridyldisulfide (3 mmoles, 10 equiv., 607 mg) was then subsequently added. The reaction was allowed to stir for 4 h. The crude product was obtained, as an off-white precipitate, by adding the reaction mixture slowly into an ice-cold solution of 100 mL acetone, 100 mL diethyl ether, and 1.25 mL of saturated sodium perchlorate ($NaClO_4$) in acetone. The precipitate was then collected by suction filtration, followed by volatile removal under high vacuum. The resulting crude residue was purified. Reverse-phase flash chromatography was carried out on a Teledyne Isco CombiFlash Rf system (Lincoln, NE), on a 50 g C18Aq column over 15 CV of 0–15% $CH_3CN$ in TEAB (pH 7.5). The titled compound was eluted from the column at ~9% $CH_3CN$ in TEAB, followed by lyophilization to afford the titled compound (in tetraethylammonium cation form) as a white fluffy solid.

**(α-L-threofuranosyl)guanine 3′-*O*-phosphoro-2-aminoimidazolide (2AIptG).** $^{1}H$ NMR (400 MHz, Deuterium Oxide) δ 7.87 (s, 1H), 6.50 (t, *J* = 2.2 Hz, 1H), 6.43 (t, *J* = 1.9 Hz, 1H), 5.89 (s, 1H), 4.86 (d, *J* = 8.4 Hz, 1H), 4.47 – 4.39 (m, 3H). $^{31}P$ NMR (162 MHz, Deuterium Oxide) δ -13.47.

**(α-L-threofuranosyl)cytidine 3′-*O*-phosphoro-2-aminoimidazolide (2AIptC).** $^{1}H$ NMR (400 MHz, Deuterium Oxide) δ 7.67 (d, *J* = 7.6 Hz, 1H), 6.72 (t, *J* = 2.3 Hz, 1H), 6.62 (t, *J* = 2.2 Hz, 1H), 5.93 (d, *J* = 7.6 Hz, 1H), 5.71 (s, 1H), 4.53 – 4.39 (m, 3H), 4.12 (s, 1H). $^{31}P$ NMR (162 MHz, Deuterium Oxide) δ -13.84.

### 2. Supplementary Figures

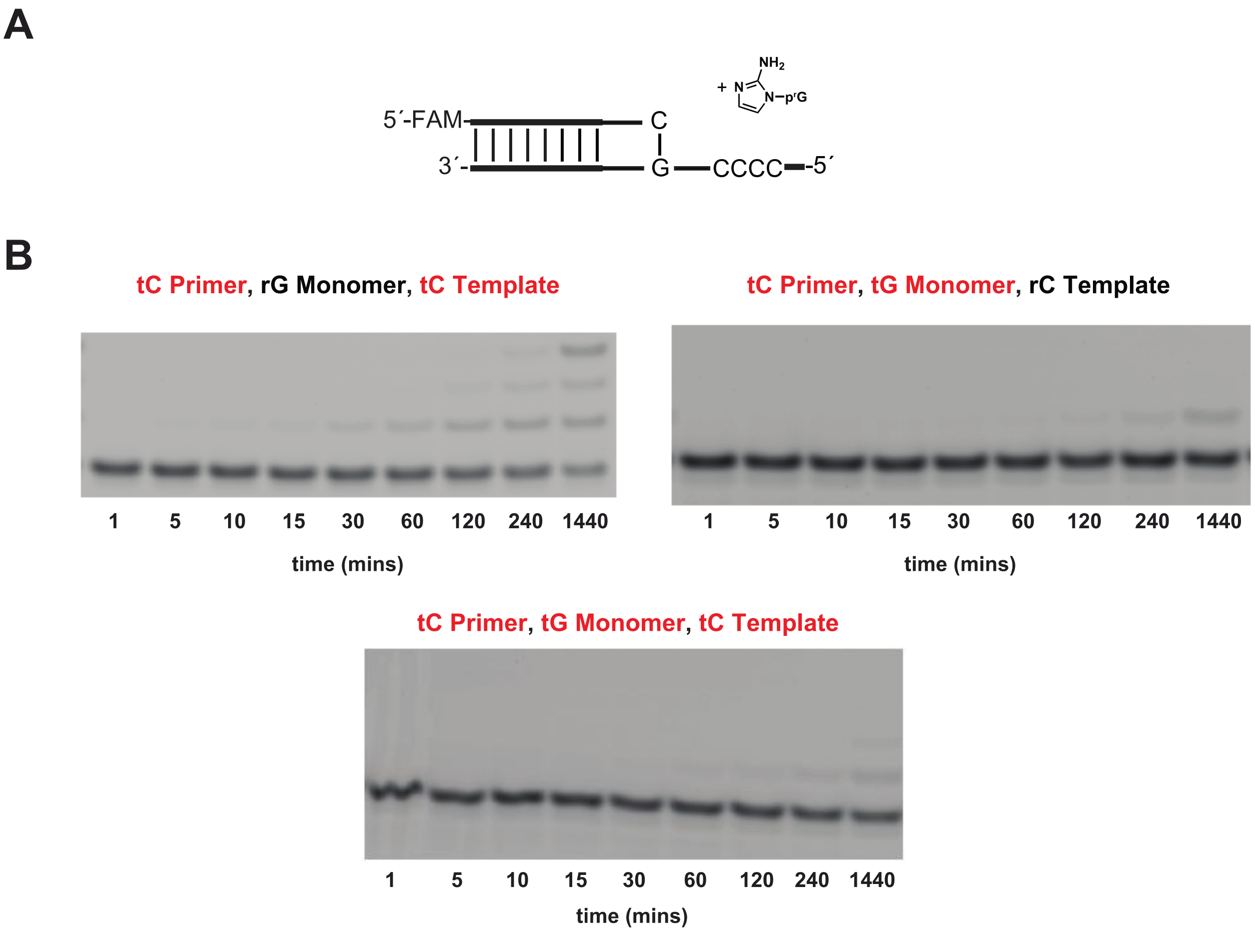


**Figure S-1.** Evaluation of nonenzymatic primer extension with TNA residues across various positions. (a) Schematic representation of a primer extension reaction with 2-aminoimidazole activated monomers (either 2AIrG or 2AItG), a primer containing either a ribo- or threo-C at the 3'-end, and a template containing either a ribo-C (rC) or threo-C (tC). (b) Gel electrophoresis images of primer extension across various conditions. All reactions were performed at pH 8.0, in 200 mM HEPES, 200 mM $Mg^{2+}$, 40 mM 2AIpG.

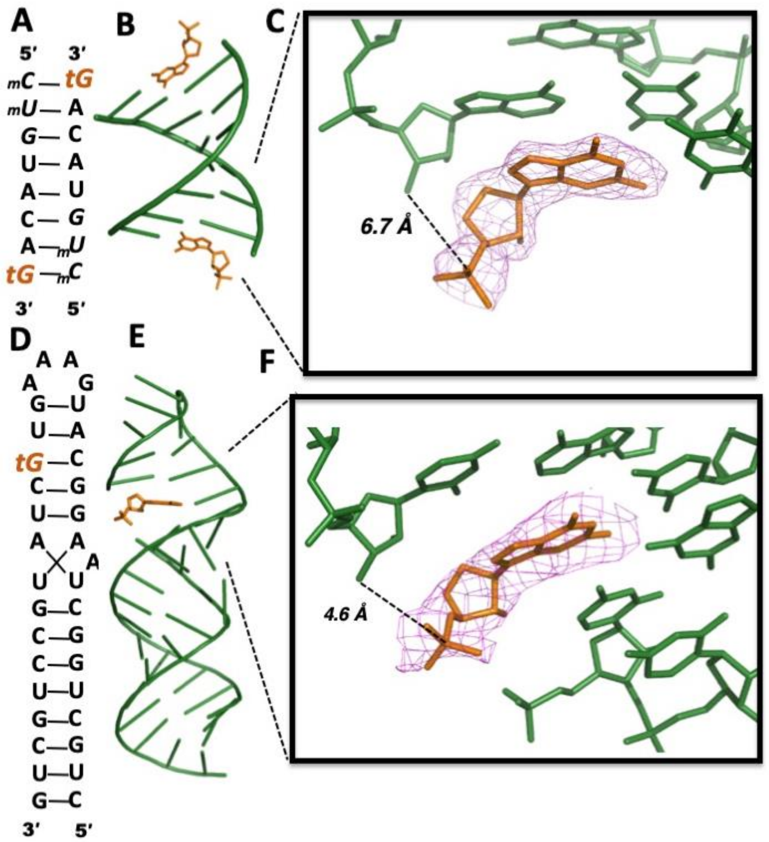


**Figure S-2**. Crystal structures of two RNA primer/template complexes with tGMP monomers. (A) Diagram and designed secondary structure of RNA primer/template/tGMP complex TNA-M1. tGMP monomers (orange) are bound at both ends of the duplex. (B) Overall crystal structure of TNA-M1 complex. (C) Local view of the tGMP bound to template by Watson-Crick base-pairing. (D) Diagram and designed secondary structure of RNA hairpin-tGMP complex TNA-M2. tGMP monomers (orange) are sandwiched between the 3′-end of the RNA primer and a downstream helper oligonucleotide. (E) Overall crystal structure of TNA-M2 complex. (F) Local view of the tGMP bound to template. Pink mesh indicates the corresponding $2F_o$-$F_c$ maps contoured at 2.0 σ and 1.5 σ. Distances between 3′-OH of the primers and the P atoms of the bound tGMPs are labelled.

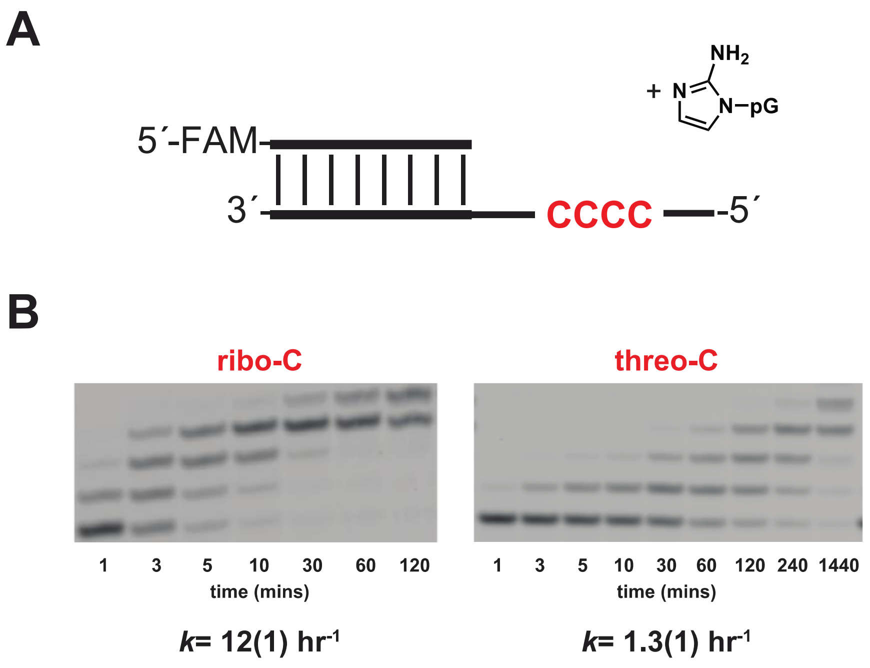


**Figure S-3.** Comparison of nonenzymatic primer extension with 2-aminoimidazole activated guanosine (ribo-G) across from four templating ribo- or threo-cytidine residues. (a) Schematic representation of a primer extension reaction of 2AIpG with template containing either ribo-C or threo-C. (b) Gel electrophoresis images and rates of primer extension for 2-aminoimidazole activated ribo-guanosine. All reactions were performed at pH 8.0, in 200 mM HEPES, 200 mM $Mg^{2+}$, 40 mM 2AIpG. Values are the mean rate constant ± SD in parentheses, with the last digit reported being the last significant figure and the one in which error arises from triplicate experiments.

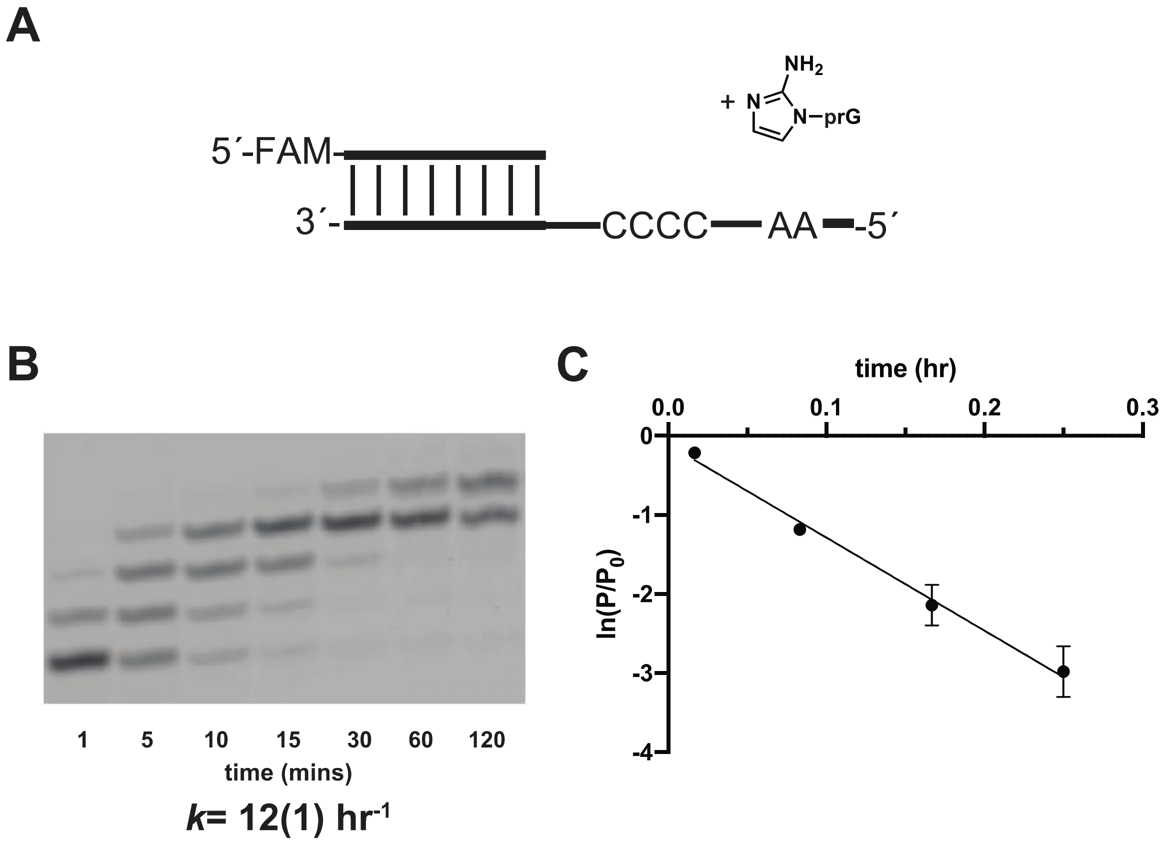


**Figure S-4.** All RNA primer extension reaction with 2-AI activated ribo-guanosine. (a) Schematic representation of a primer extension reaction. (b) Representative PAGE analysis of result. (c) Plot of $\ln(P/P_0)$ as a function of time. The rate of extension was determined from linear least-square fits of the data from three independent experiments. All reactions were carried out in triplicate using 40 mM 2AIpG, pH 8.0, in 200 mM HEPES, 200 mM $Mg^{2+}$.

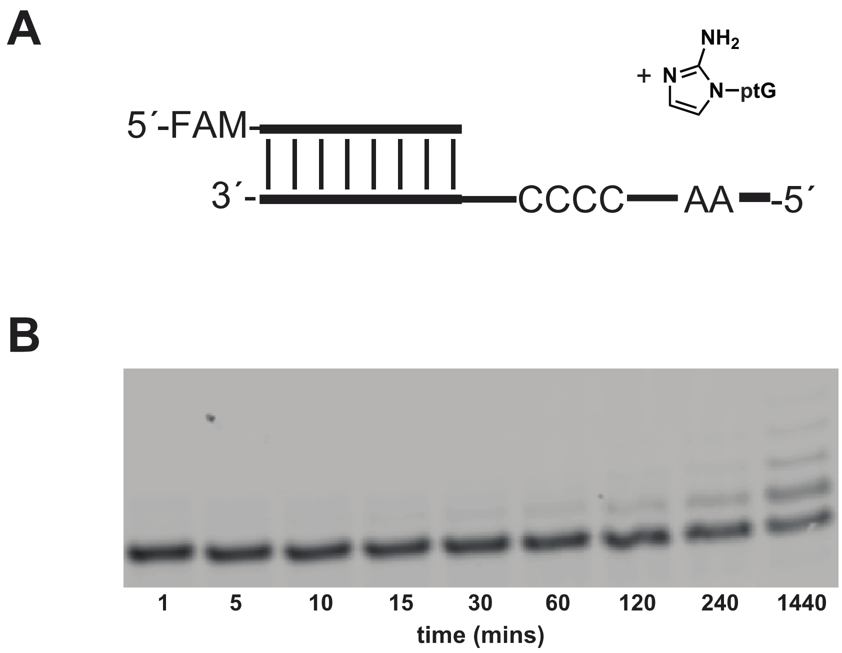


***k*< 0.1 hr$^{-1}$**

**Figure S-5.** Primer extension with 2AI-activated threo-guanosine monomer, RNA primer and RNA template. (a) Schematic representation of a primer extension reaction. (b) Representative PAGE analysis of result. (c) Plot of ln($P/P_0$) as a function of time. The rate of extension was determined from linear least-square fits of the data from three independent experiments. All reactions were carried out in triplicate using 40 mM 2AIptG, pH 8.0, in 200 mM HEPES, 200 mM $Mg^{2+}$.

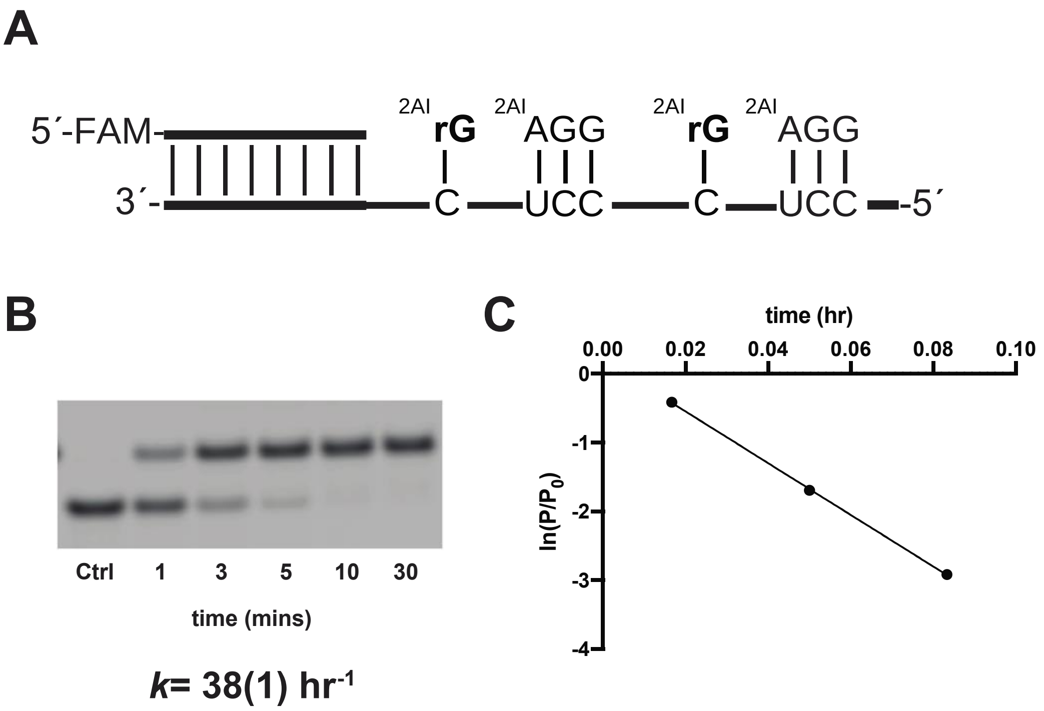


**Figure S-6.** Control all-RNA primer extension reaction. (a) Schematic representation of a primer extension reaction. (b) Representative PAGE analysis of result. (c) Plot of $\ln(P/P_0)$ as a function of time. The rate of extension was determined from linear least-square fits of the data from three independent experiments. All reactions were carried out in triplicate using 20 mM 2AIprG, 1 mM 2AIpAGG, pH 8.0, in 200 mM HEPES, 200 mM $Mg^{2+}$.

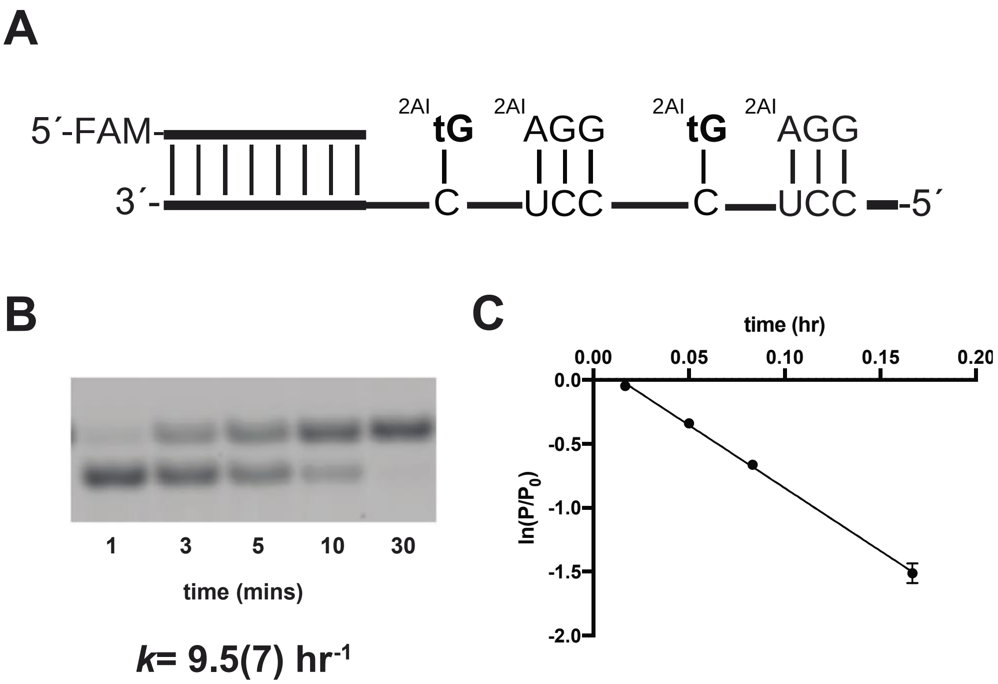


**Figure S-7.** Primer extension reaction with 2AI-activated threo-guanosine, with RNA primer, template, and activated helper. (a) Schematic representation of a primer extension reaction. (b) Representative PAGE analysis of result. (c) Plot of ln($P/P_0$) as a function of time. The rate of extension was determined from linear least-square fits of the data from three independent experiments. All reactions were carried out in triplicate using 20 mM 2AIptG, 1 mM 2AIpAGG, pH 8.0, in 200 mM HEPES, 200 mM $Mg^{2+}$.

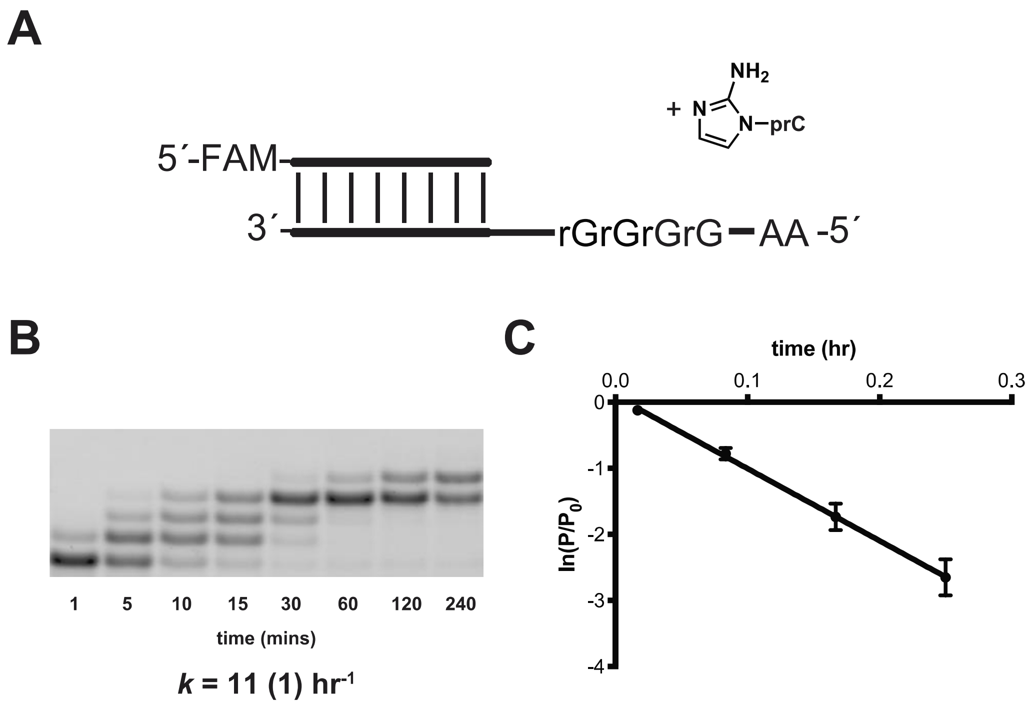


**Figure S-8.** Primer extension reaction 2AI-activated ribo-C monomer, and RNA primer and template. (a) Schematic representation of a primer extension reaction. (b) Representative PAGE analysis of result. (c) Plot of ln($P/P_0$) as a function of time. The rate of extension was determined from linear least-square fits of the data from three independent experiments. All reactions were carried out in triplicate using 40 mM 2AIpC, pH 8.0, in 200 mM HEPES, 200 mM $Mg^{2+}$.

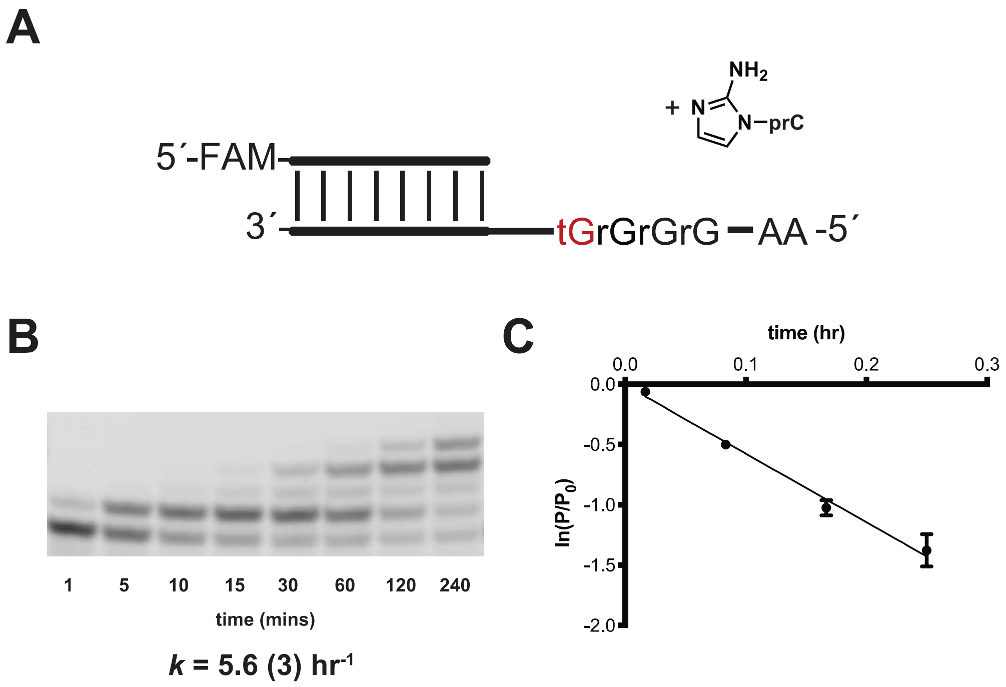


**Figure S-9.** Primer extension across from a single threo-nucleotide in the RNA template. (a) Schematic representation of the primer extension reaction, showing the RNA primer and monomer. (b) Representative PAGE analysis of result. (c) Plot of $\ln(P/P_0)$ as a function of time. The rate of extension was determined from linear least-square fits of the data from three independent experiments. All reactions were carried out in triplicate using 40 mM 2AIprC, pH 8.0, in 200 mM HEPES, 200 mM $Mg^{2+}$.

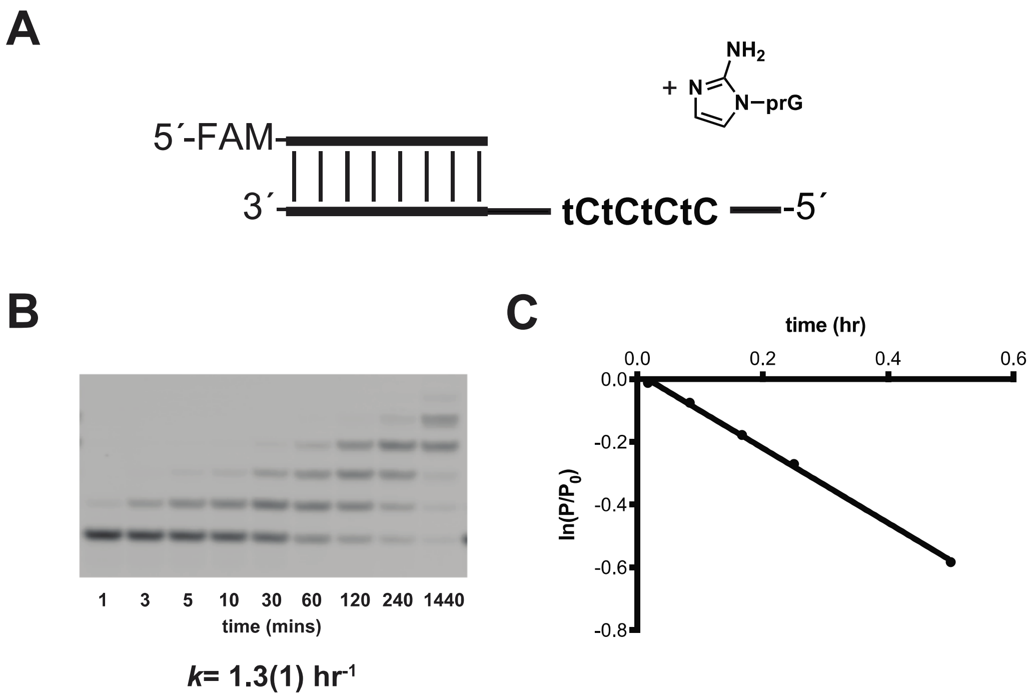


**Figure S-10.** Primer extension across a template region consisting of four threo-C residues. (a) Schematic representation of the primer extension reaction, showing the RNA primer and the 2AI-activated ribo-guanosine monomer. (b) Representative PAGE analysis of result. (c) Plot of $\ln(P/P_0)$ as a function of time. The rate of extension was determined from linear least-square fits of the data from three independent experiments. All reactions were carried out in triplicate using 40 mM 2AIpG, pH 8.0, in 200 mM HEPES, 200 mM $Mg^{2+}$.

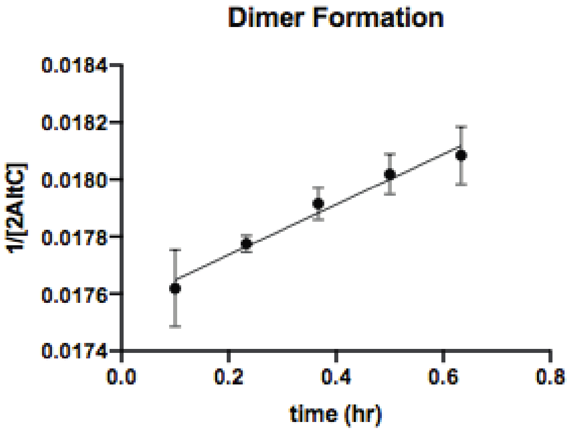


$k = 8.8 \pm 0.5 * 10^{-4}$ $mM^{-1}$ $hr^{-1}$

**Figure S-11**. Kinetic analysis of imidazolium-bridged tC dimer (tC*tC) formation. All reactions were carried out in triplicate using 60 mM activated threo-cytidine (2AIptC), 200 mM $Na^{+}$-HEPES pH 8.0, and 10% $D_2O$. As standard for second-order kinetics, a linearized plot of 1/[2AIptC] as a function of time is shown above and the rate of formation was determined from linear least-square fits of the data from an average of three independent experiments.

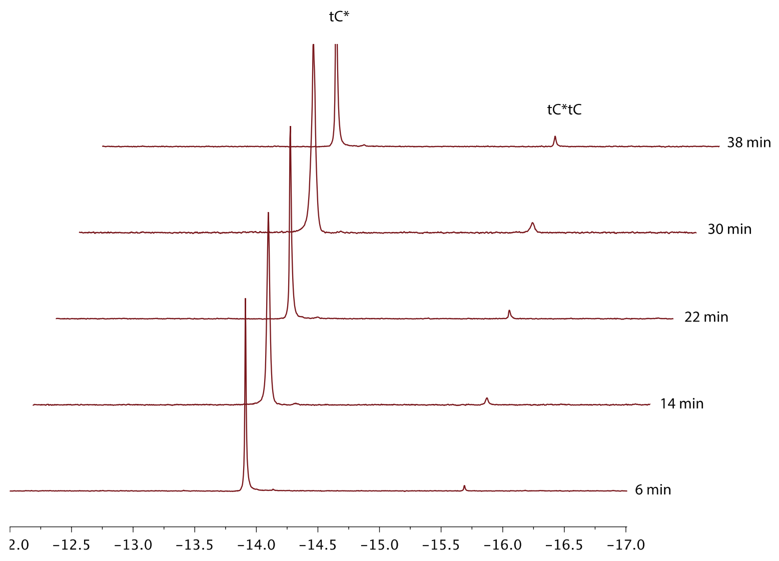


**Figure S-12.** $^{31}$P NMR spectrum of the formation of tC imidazolium-bridged dimer tC*tC (60 mM of activated monomer 2AIptC (tC*), 200 mM $Na^+$-HEPES pH 8.0, and 10% $D_2O$) after 6, 14, 22, 30, and 38 minutes of reaction time (bottom to top).

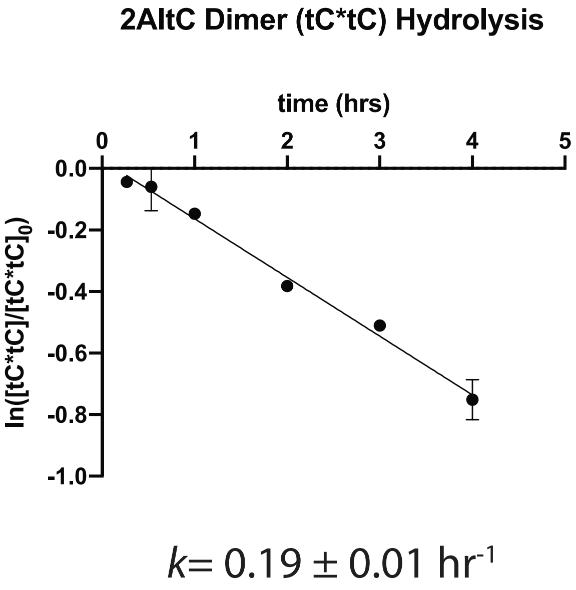


**Figure S-13**. Kinetic analysis of the hydrolysis of imidazolium-bridged tC dimer (tC*tC). The reaction was carried out in triplicate using 5 mM activated threo-cytidine species, 50 mM $MgCl_2$, 200 mM $Na^+$-HEPES pH 8.0, and 10% $D_2O$. A linearized plot of ln([tC*tC]/[tC*tC]$_0$) as a function of time is shown above and the rate of hydrolysis was determined from linear least-square fits of the data from an average of three independent experiments.

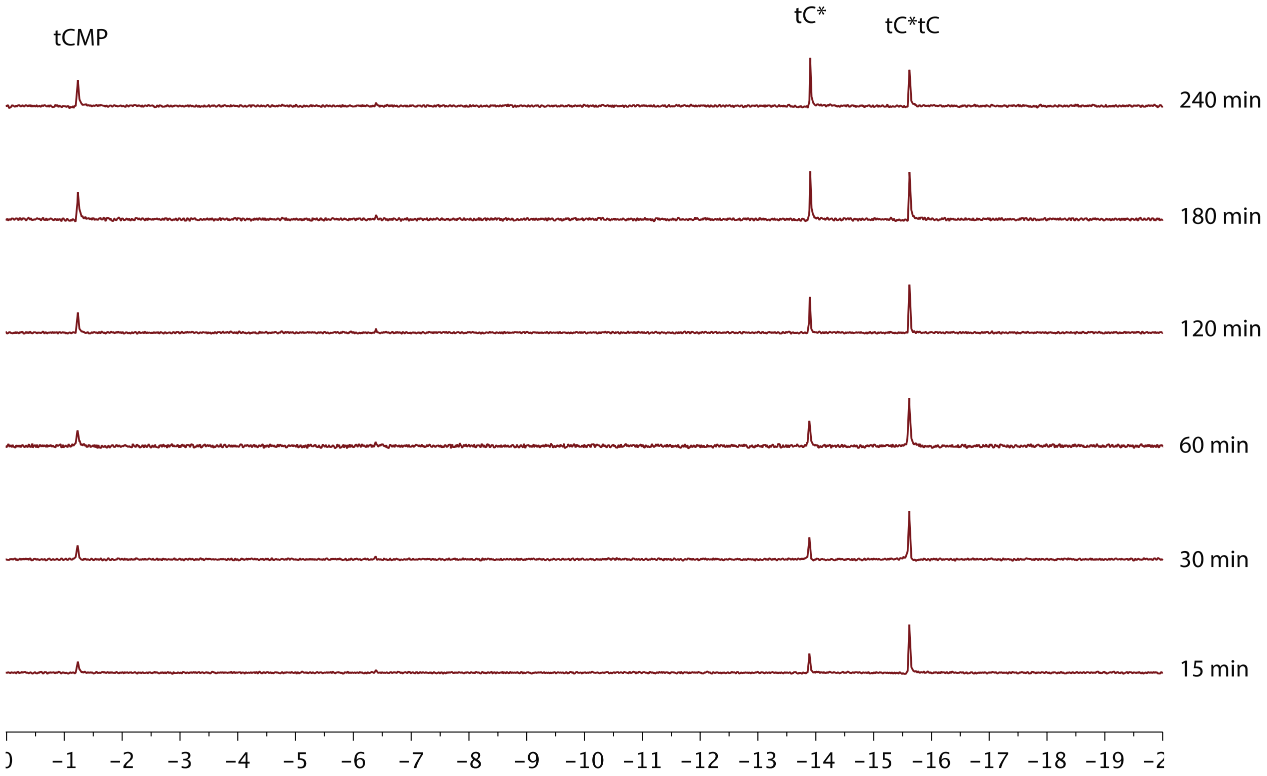


**Figure S-14.** $^{31}$P NMR spectrum of the hydrolysis of tC imidazolium-bridged dimer under nonenzymatic primer extension condition (5 mM of activated species, 50 mM $MgCl_2$, 200 mM $Na^+$-HEPES pH 8.0, and 10% $D_2O$) after 15, 30, 60, 120, 180, and 240 minutes of reaction time (bottom to top).

### 3. X-ray Crystallographic Studies.

Optimal crystallization conditions, data collection, phasing, and refinement statistics of the determined structures are listed in Tables S1, S2 and S3.

**Table S1. Optimized conditions for crystallization of RNA-monomer complexes**

| | Optimized crystallization conditions |
|---|---|
| TNA-T1 | 0.2 M Lithium sulfate monohydrate, 0.1 M HEPES pH 7.5, 25% w/v Polyethylene glycol 3,350 |
| TNA-P1 | 3.5 M Sodium formate pH 7.0 |
| TNA-M1 | 25 % v/v Polyethylene glycol 400, 100 mM tri-Sodium citrate; pH 5.6 130 mM Sodium chloride, 60 mM Magnesium chloride |
| TNA-M2 | 1 M Lithium sulfate, 50 mM tri-Sodium citrate, 3 % w/v 2-Propanol, 50 mM HEPES; pH 7.5 |
| TNA-D1 | 10 % w/v Polyethylene glycol 6,000, 50 mM HEPES; pH 7.0, 200 mM Ammonium acetate, 150 mM Magnesium acetate |

**Table S2. Data collection statistics.**

| **Structure** | TNA-T1 | TNA-P1 |
|---|---|---|
| Space group | R3 | R3 |
| Unit cell parameters (Å, º) | 68.77, 68.77, 69.96, 90, 90, 120 | 71.32, 71.32, 67.57, 90, 90, 120 |
| Resolution range, Å (last shell) | 50-2.95 (3.06-2.95) | 50-2.7 (2.8-2.7) |
| Unique reflections | 2599 | 3369 |
| Completeness, % | 98.5 (89.6) | 97.5 (89.4) |
| $R_{merge}$, % | 0.116 (0.332) | 0.117 (0.415) |
| <I/σ(I)> | 10.3 (1.83) | 9.5 (2.3) |
| Redundancy | 6.0 (4.5) | 3.5 (2.8) |

| **Structure** | TNA-M1 | TNA-M2 | TNA-D1 |
|---|---|---|---|
| Space group | P63 | R3 | P3121 |
| Unit cell parameters (Å, º) | 51.89, 51.89, 37.89, 90.00, 90.00, 120.00 | 69.63, 69.63, 70.51, 90.00, 90.00, 120.00 | 43.96, 43.96, 84.52, 90.00, 90.00, 120.00 |
| Resolution range, Å | 50-2.36 (2.44-2.36) | 50-2.80 (2.90-2.80) | 50-2.4 (2.49-2.4) |

| | | | |
|---|---|---|---|
| (last shell) | | | |
| Unique reflections | 2444 | 3110 | 4009 |
| Completeness, % | 99.9 (99.6) | 99.5 (96) | 99.9 (100) |
| $R_{merge}$, % | 0.163 (0.537) | 0.10 (0.567) | 5.9 (70.9) |
| <I/σ(I)> | 10.7 (1.89) | 15.7 (1.42) | 37.2 (3.1) |
| Redundancy | 4.5 (3.9) | 5.4 (4.1) | 10.2 (10.6) |

### Table S3. Structure refinement statistics.

| **Structure** | TNA-T1 | TNA-P1 |
|---|---|---|
| PDB code | 6U7Y | 6U7Z |
| RNA duplex per asymmetric unit | 1 | 1 |
| Resolution range, Å | 45.3-2.95 | 45.6-2.71 |
| $R_{work}$, % | 18.2 | 17.1 |
| $R_{free}$, % | 25.6 | 23.4 |
| Number of reflections | 2463 | 3207 |
| Bond length R.M.S. (Å) | 0.008 | 0.007 |
| Bond angle R.M.S. | 1.75 | 1.63 |
| Average B-factors, (Å$^2$) | 62.6 | 87.6 |

| **Structure** | TNA-M1 | TNA-M2 | TNA-D1 |
|---|---|---|---|
| PDB code | 6U89 | 6U8F | 6U8U |
| RNA duplex per asymmetric unit | 1 | 1 | 1 |
| Resolution range, Å | 44.96-2.36 | 45.83-2.81 | 50-2.4 |
| $R_{work}$, % | 18.8 | 17.5 | 21.6 |
| $R_{free}$, % | 27.4 | 20.9 | 26.7 |
| Number of reflections | 2332 | 2966 | 3789 |
| Bond length R.M.S. (Å) | 0.02 | 0.01 | 0.028 |
| Bond angle R.M.S. | 2.96 | 1.95 | 2.92 |
| Average B-factors, (Å$^2$) | 36.2 | 81.1 | 53.9 |
